# Validation of carbon isotopologue distribution measurements by GC-MS and application to ^13^C-metabolic flux analysis of the tricarboxylic acid cycle in *Brassica napus* leaves

**DOI:** 10.1101/2022.03.03.482145

**Authors:** Younès Dellero, Solenne Berardocco, Cécilia Berges, Olivier Filangi, Alain Bouchereau

## Abstract

The estimation of metabolic fluxes in photosynthetic organisms represents an important challenge that has gained interest over the last decade with the development of ^13^C-Metabolic Flux Analysis at isotopically non-stationary steady-state. This approach requires a high level of accuracy for the measurement of Carbon Isotopologue Distribution in plant metabolites. But this accuracy has still not been evaluated at the isotopologue level for GC-MS, leading to uncertainties for the metabolic fluxes calculated based on these fragments. Here, we developed a workflow to validate the measurements of CIDs from plant metabolites with GC-MS by producing tailor-made *E. coli* standard extracts harboring a predictable binomial CID for some organic and amino acids. Overall, most of our TMS-derivatives mass fragments were validated with these standards and at natural isotope abundance in plant matrices. Then, we applied this validated MS method to investigate the light/dark regulation of plant TCA cycle by incorporating U-^13^C-pyruvate to *Brassica napus* leaf discs. We took advantage of pathway-specific isotopologues/isotopomers observed between two and six hours of labeling to show that the TCA cycle can operate in a cyclic manner under both light and dark conditions. Interestingly, this forward cyclic flux mode has a nearly four-fold higher contribution for pyruvate-to-citrate and pyruvate-to-malate fluxes than the phosphoenolpyruvate carboxylase (PEPc) flux reassimilating carbon derived from some mitochondrial enzymes. The contribution of stored citrate to the mitochondrial TCA cycle activity was also questioned based on dynamics of ^13^C-enrichment in citrate, glutamate and succinate and variations of citrate total amounts under light and dark conditions. Interestingly, there was a light-dependent ^13^C-incorporation into glycine and serine showing that decarboxylations from pyruvate dehydrogenase complex and TCA cycle enzymes were actively reassimilated and could represent up to 5% to net photosynthesis.

## 1 INTRODUCTION

Metabolic fluxes within central metabolism are among the most important determinants of plant cell physiology, supporting central energetical functions and allowing them to quickly adapt to environmental changes by controlling metabolite turnover (Clark et al., 2020). They can be estimated from ^13^C-Metabolic Flux Analysis (MFA), which relies on the tracing of ^13^C throughout a metabolic network in a relative metabolic steady-state to model and/or compare metabolic fluxes (Ratcliffe and Shachar-Hill, 2006;Allen and Young, 2019;Antoniewicz, 2021). In phototrophic tissues, ^13^C-MFA can only be performed at isotopically non-stationary steady-state (INST-MFA) and requires a high level of accuracy for the measurement of ^13^C enrichment within metabolites (Cheah and Young, 2018;Cheah et al., 2020;Antoniewicz, 2021;Treves et al., 2022). Since small errors in mass isotopologue distribution can propagate to large errors in estimated fluxes (Antoniewicz et al., 2006), an accurate measurement of ^13^C-enrichment still remains an important prerequisite to explore plant metabolism dynamics through isotopic labeling approaches. Two major analytical methods are currently employed to analyze the Carbon Isotope Distribution (CID) within plant metabolites: Nuclear Magnetic Resonance (NMR) and Mass Spectrometry (MS). These methods can provide information either at the isotopologue level, *i.e*. there is a separation of labeled molecules according to their numbers of ^13^C atoms for a given metabolite, or at the isotopomer level, *i.e*. there is a separation of labeled molecules according to the position of ^13^C atoms for a given metabolite (Des Rosiers et al., 2004;Ratcliffe and Shachar-Hill, 2006). While NMR analysis of the ^13^C enrichment can provide positional labeling information, it has not been widely used in plants due to the need of large amounts of fresh material to overcome its low sensitivity at natural ^13^C abundance, thus preventing the possibility to perform time-course ^13^C-isotope tracing experiments (Kruger et al., 2007;Masakapalli et al., 2010;Dellero et al., 2016;Abadie et al., 2017;Abadie and Tcherkez, 2021). Conversely, mass spectrometry has been preferentially used to analyze the Carbon Isotope Distribution (CID) within plant metabolites during time-course ^13^C-isotope tracing experiments (Szecowka et al., 2013;Ma et al., 2014;Medeiros et al., 2018;Treves et al., 2022). This method is highly sensitive at natural ^13^C abundance but essentially gives information at the isotopologue level, given that MS fragments can hardly cover the complete set of all possible isotopomers for a given metabolite (Lima et al., 2021). Interestingly, the recent analysis of CID for amino acids by LC-MS revealed some biases and errors in the measurement of CID (Heuillet et al., 2018), thus highlighting the need to also evaluate measurement accuracy of MS methods for subsequent use in ^13^C-MFA of plants.

Among plant metabolites, organic and amino acids are major players for plant primary metabolism. They are actively involved in crucial physiological processes such as photorespiration, *de novo* nitrogen assimilation and mitochondrial respiration (Xu et al., 2012;Zhang and Fernie, 2018;Dellero, 2020;Timm, 2020). Interestingly, most of the carbon backbones for amino acid biosynthesis come from organic acids of the tricarboxylic acid (TCA) cycle, raising this cycle as a central hub for plant primary metabolism (Araujo et al., 2012;Galili et al., 2016;Tcherkez et al., 2017). The functioning and regulation of the plant TCA cycle has been described based on flux balance models (Sweetlove et al., 2010;Araujo et al., 2012;Cheung et al., 2014) but also based on ^13^C-labeling studies with different cell types/experimental setups: attached leaves (Xu et al., 2021;Xu et al., 2022), detached leaves (Tcherkez et al., 2009;Gauthier et al., 2010;Zhang et al., 2021), leaf discs (Le et al., 2021), guard cell cultures (Daloso et al., 2017), heterotrophic cell cultures (Masakapalli et al., 2010;Masakapalli et al., 2014) and isolated mitochondria (Le et al., 2021;Le et al., 2022). Overall, most of these works proposed that the plant TCA cycle is not cyclic in the light, with two branches acting separately: i) a first branch stimulated by the anaplerotic activity of phosphoenolpyruvate carboxylase (PEPc) leading to the production of oxaloacetate, malate and fumarate; ii) a second branch stimulated by the entry of citrate leading to the production of isocitrate, 2-oxoglutarate, glutamate and sometimes succinate. These conclusions were notably supported by the inhibition of the pyruvate dehydrogenase complex (PDC) and the mitochondrial respiration in the light while nitrogen assimilation was relatively high compared to dark conditions and required important amounts of 2-oxoglutarate (Tcherkez et al., 2005;Gauthier et al., 2010). In addition, the photorespiratory activity of the mitochondrial glycine decarboxylase complex could largely compete with the second branch of the TCA cycle for NADH production, thereby sustaining most of the mitochondrial energetic production in the light, independently of the activity of TCA cycle enzymes (Timm and Hagemann, 2020). Besides this, the requirement to support continued export of sugar and amino acids from the leaf during the night and to meet cellular maintenance costs was also invoked to explain the light-dependent bi-directional functioning of the TCA cycle (Cheung et al., 2014). Regarding the metabolic origin of citrate for the second branch in the light, it can come from: i) TCA cycle *de novo* biosynthesis using acetyl-CoA derived from either glycolysis and alanine transamination; ii) a stored citrate pool in a light/dark dependent-manner; iii) the peroxisomal glyoxylate cycle (Gauthier et al., 2010;Sweetlove et al., 2010;Cheung et al., 2014;Dong et al., 2018;Le et al., 2021). Based on the NMR analysis of combined ^13^CO_2_ and ^15^NH_4_^15^NO_3_ labeling experiments in *Brassica napus* leaves, it was proposed that a high proportion of stored and non-labeled citrate (vacuolar pool probably) was involved in the synthesis of 2-oxoglutarate under light conditions to sustain *de novo* nitrogen assimilation through glutamate (Gauthier et al., 2010). Under this scenario, the citrate pool would be replenished during dark conditions from glycolysis-derived ^13^C-pyruvate, leading to a higher ^13^C enrichment of the citrate after a light/dark cycle. Thus, the light and dark environmental conditions are expected to introduce important variations of metabolic fluxes within the plant TCA cycle (cyclic versus non-cyclic) and its associated PEPc anaplerotic pathway in *B. napus* leaves. Nevertheless, these variations have still not been carefully investigated with time-course ^13^C-isotope tracing experiments and pool size quantification/estimation (^13^C-MFA concepts), thus casting doubt on previous conclusions drawn in *B. napus* from NMR-based experiments (one time-point, no pool size quantification/estimation). In addition, recent ^13^CO_2_ labeling experiments coupled to ^13^C-INST-MFA succeeded to model leaf primary metabolism (including TCA cycle) without the need of a stored ^12^C Citrate pool in *Camelina sativa* and Arabidopsis leaves (Xu et al., 2021;Xu et al., 2022).

To date, the routine analysis of CID for plant organic and amino acids has been often assessed by GC-MS using TMS-derivatives (Szecowka et al., 2013;Araujo et al., 2014;Lima et al., 2018;Medeiros et al., 2018;Lima et al., 2021;Treves et al., 2022) although previous comparisons of two derivatization methods for GC-MS suggested that TBDMS-derivatives were potentially less prone to errors for amino acid analysis than TMS-derivatives (Antoniewicz et al., 2007;Young et al., 2014). However, the suitability of molecular ions from TMS and TBDMS derivatives was only performed using U-^13^C-metabolites in each of these studies, which did not evaluate the entire CID but only the carbon distribution between the unlabeled isotopologue and the fully labeled isotopologue (M0 and M5 for a 5-carbon metabolite for example). Given that isotopic effects and analytical biases may occur for any isotopologue of each tested metabolite, it is essential to validate MS measurements for each isotopologue. This validation step is even more important for plants, given that plant metabolites show strong differences of ^13^C-enrichment dynamics between amino and organic acids (Szecowka et al., 2013). For this purpose, the recent development of an *E. coli* ^13^C biological standard with controlled patterns of CID for organic and amino acids may help to address this issue (Millard et al., 2014). Indeed, the recent use of these reference samples allowed to identify important analytical biases in the CID measurement of Leucine and Asparagine in a concentration-dependent manner by LC-MS (Heuillet et al., 2018). Interestingly, this validated LC-MS method successfully measured nitrogen isotope distribution within plant amino acids and led to the estimation of metabolic fluxes from ^15^N-isotope transient labeling experiments in *B. napus* leaves at a metabolic steady-state using the ScalaFlux approach (Dellero et al., 2020b;Heuillet et al., 2020;Millard et al., 2020).

Here, we proposed a workflow to assess the measurement accuracy of CID from organic and amino acids by GC-MS for applications in plants. This methodology was based on chemical standards, plant samples at natural ^13^C abundance and two specific *E. coli* ^13^C reference samples (^13^C-PT), produced to specifically harbor controlled patterns of CID for organic and amino acids (Pascal’s Triangle coefficients). Next, we applied our validated MS method to investigate the light/dark regulation of metabolic fluxes within plant TCA cycle by performing U-^13^C-pyruvate labeling kinetic experiments on *B. napus* leaf discs under light or dark conditions. Using ^13^C-MFA principles and some specific isotopologues, we questioned the contribution of stored citrate to the TCA cycle activity and showed that the TCA cycle partially works as a cycle under light conditions in *B. napus*. We also analyzed the fractional contribution of phosphoenolpyruvate carboxylase (PEPc) and TCA cycle for pyruvate-to-malate and for pyruvate-to-citrate fluxes.

## 2 MATERIALS AND METHODS

### Organisms and culture

The strain *Escherichia coli* K-12 MG1655 was grown on minimal medium containing 5 mM KH_2_PO_4_,10mM Na_2_HPO_4_, 9 mM NaCl, 40 mM NH_4_Cl, 0.8 mM MgSO_4_, 0.1 mM CaCl_2_, 0.3 mM thiamine, and 45 mM of the ^13^C-labeled acetate mixture as the sole carbon source. Acetate and thiamine were sterilized by filtration, whereas the other compounds were autoclaved separately. Batch cultures were carried out in 1-L baffled flasks with 500 mL of medium sparged with synthetic gas (80% N_2_/20%O_2_ containing <1 ppm CO_2_, Air Liquide). Cultivation was performed in a Multifors Bioreactor (Infors HT, Bottmingen-Basel, Switzerland) at 37 °C and 200 rpm of orbital shaking. The pH was maintained at 7.0 by the addition of 2 M HCl. Cell growth was monitored by measuring optical density at 600 nm with a Genesys 6 spectrophotometer (Thermo, Carlsbad, CA, USA). Amount of biomass in each sample was calculated using a coefficient of 0.37 g of cell dry weight per OD unit. Arabidopsis thaliana seeds were sterilized for 10 min with absolute ethanol containing 7.5% NaClO and rinsed two times with absolute ethanol. Seeds were sowed on ½ MS media containing 1% Agar and incubated to the dark at 4°C for 48-72 hours to break seed dormancy. Seedlings were grown at ambient air under a day/night cycle with 16 h light/8h dark at a light intensity of 120 μmol photons.m^−2^.s^−1^ and 20°C day/18°C night. Seedlings were harvested after 2-3 weeks by instant sampling with liquid nitrogen and lyophilized for 96 h before to be stored at −80°C until analysis. Two-month old *Brassica napus* plants were obtained from seeds exactly as described before (Dellero et al., 2020a). The leaf L9 was used for all experiments (natural ^13^C abundance and U-^13^C-pyruvate incorporation).

### U-^13^C-pyruvate labeling experiments

Four hours after the beginning of the light period, each leaf was cut at the basis of the petiole and then 66 leaf discs (0.8 cm^2^) were randomly punched with a cork-borer in both laminas of each leaf. Leaf discs were first floated in a 10 mM MES-KOH (pH 6.5) buffer for 15 min in order to process the L9 leaf for four plants (see details for leaf annotation in (Dellero et al., 2020a)). In this experiment, one leaf of a plant corresponded to a biological replicate. Then for each time point, six leaf discs from the same leaf were transferred in a well of a 6-well microplate filled with 4 mL/well of a buffer containing 10 mM MES-KOH (pH 6.5) and 10 mM U-^13^C-pyruvate (^13^C isotopic purity of 99%; sodium salt from Cambridge Isotope Laboratories (Eurisotop, France)). All incubations were performed at 20 °C with an orbital shaking of 70 rpm under either a continuous light of 120 μmol photons m^−2^ s^−1^ or under dark conditions using multiple opaque plastic bags, similarly to the previous experimental set-up used for dark-induced senescence (Dellero et al., 2020a). The timepoint 0 was the same for both conditions as it corresponded the beginning of the experiment. For each time point (0, 0.5, 1, 2, 4, 6 hours of incubation), the leaf discs were harvested and rinsed three times (10 sec) with a neutral buffer (i.e., without U-^13^C-pyruvate) and dried for 10 s on a clean tissue before being frozen in liquid nitrogen and stored at −80 °C. The timepoint 0 of the experiment was incubated for 2 sec with the labeled buffer before being directly rinsed. This was done to evaluate the residual amount of ^13^C_3_-pyruvate detected in our analysis that was not truly incorporated into leaf discs (a mean ^13^C enrichment close to 35% for pyruvate at T0 was observed in our study). This amount reflected probably biases in the washing procedure and/or an artificial fixation of ^13^C_3_-pyruvate to the leaf disc surface.

### Analysis of ^13^C-labeled Acetate by NMR

An equimolar mixture of all ^13^C-isotopomers of acetate was produced from ^12^C-acetate, 1-^13^C-acetate, 2-^13^C-acetate and U-^13^C-acetate (^13^C isotopic purity of 99%, sodium salts from Cambridge Isotope Laboratories (Eurisotop, France); **Figure 1**). Quantitative 1H 1D-NMR was used to control the isotopic composition of the ^13^C-labeled substrate before use. NMR spectra were recorded with an Avance 500 MHz spectrometer (Bruker, Rheinstetten, Germany) at 298 K, using a 30° pulse and a relaxation delay of 20 s. The proportion of each isotopic form was quantified by fitting using a mixed Gaussian−Lorentzian model (Millard et al., 2017). We obtained a molecular ^13^C enrichment of 49.9 % ± 0.8.

**Figure 1.**
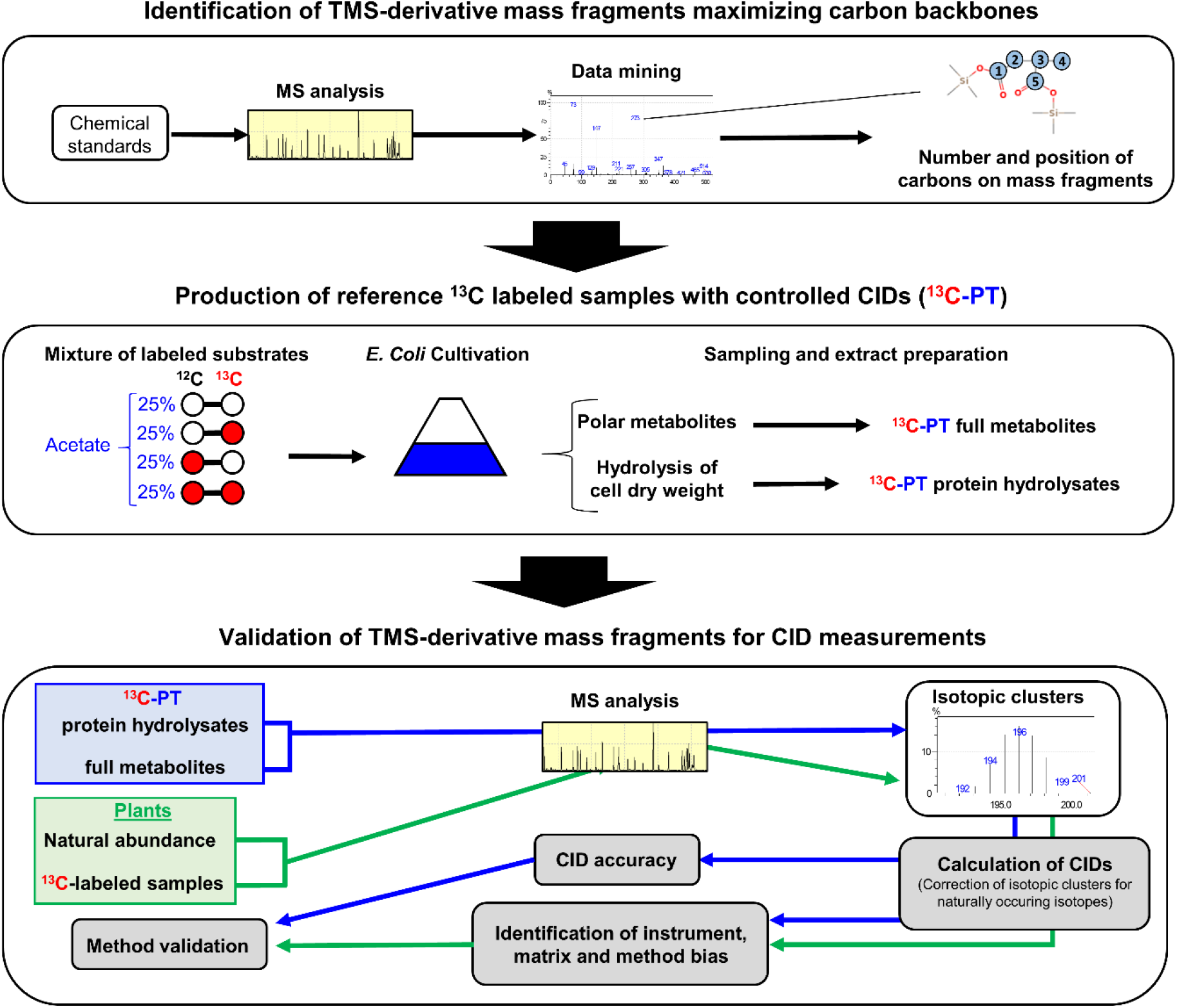
Workflow used in this study. A GC-MS method was built with abundant molecular ions deriving from TMS-derivatives of organic and amino acids that contained a large number of informative carbons. Then this method was evaluated by using: i) chemical standards and tailor-made ^13^C-standards that had a predictable CID using *E. coli* as a metabolic factory (^13^C-PT); ii) unlabeled and ^13^C-labeled plant samples.

### Metabolite extraction and quantification

^13^C-PT full metabolites and ^13^C-PT protein hydrolysates samples were obtained following procedures described previously (Heuillet et al., 2018). Briefly, for the ^13^C-PT full metabolites extract, 1 mL of *E. coli* broth at OD600 nm = 1.7 was micro-filtered and the pellet was rapidly quenched at −20 °C into a mixture of acetonitrile/methanol/H_2_0 (2:2:1) with 0.1% formic acid. After a centrifugation step (7000g, −20 °C, 15 min), the supernatant was evaporated and resuspended in 250 μL of Milli-Q water, to be stored at −80 °C until analysis. For the ^13^C-PT protein hydrolysates extract, the pellet obtained with the cellular extract was subjected to an acid hydrolysis with 6N HCl at 110 °C overnight. Samples were evaporated and rinsed 3 times with Milli-Q water. After a centrifugation step (10 min, 12 000g), the supernatant was stored at −80 °C. Polar metabolites of freeze-dried Arabidopsis seedlings and *B. napus* leaves were extracted with a MeOH/chloroform/H_2_O mixture following procedures described previously (Dellero et al., 2020b). The quantification of amino acid content was performed by HPLC-UV exactly as described previously (Dellero et al., 2020b). The quantification of organic acids was performed by GC-FID exactly as described previously (Bianchetti et al., 2021).

### Determination of Carbon Isotopologue Distribution by GC-MS

Dried aliquots of samples were derivatized on a MultiPurpose Sampler Gerstel and analyzed by GC-MS (GC-2010 Plus Shimadzu coupled to a triple quadrupole mass spectrometer GCMS-TQ8040 Shimadzu). Polar metabolites were first derivatized for 1 h at 40°C with 50 μL of pyridine containing 20 g.l^−1^ methoxyamine chloride and then were incubated with 50 μL of N-Methyl-N-(trimethylsilyl)-trifluoroacetamide (MSTFA) for 0.5 h at 40°C. 1 μL was injected with a split mode 1/20 at 260°C on a TG-5MS column (26098-1420, Thermo Scientific). Separation was performed in a Helium gas-stream at 1 mL.min^−1^ using the following temperature ramp : 4 min at 70°C, +10°C.min^−1^ until 198°C, 2 min at 198°C, +1°C.min^−1^ until 202°C, +15°C.min^−1^ until 268°C, 3 min at 268°C, +1°C.min^−1^ until 272°C,+10°C.min^−1^ until 310°C and 7 min à 310°C. The transfer line was maintained at 300°C and the ion source at 200°C. Ionization was performed by electron impact at 70 eV and the MS acquisition was carried out after a solvent delay of 5 min in full scan mode at 7 spectra.s_-1_ over the m/z range 45-700. MS Spectra were processed with the software “GCMS Solutions” supplied by Shimadzu to adjust retention times between each run using a mixture of alkanes (C7 to C40, Sigma-Aldrich). Peak identity was established according to their retention time and mass fragmentation spectra with respect to authentic standards derivatized (**Table S1**), and by reference to mass spectra of derivatives in the US National Institute of Standards and Technology database and in the Golm metabolome database. All mass isotopologue raw areas were subjected to correction for the contribution of naturally occurring isotopes of all elements except the carbon arising from the metabolite backbone using the IsoCor v2.1.4 package of Python (https://github.com/MetaSys-LISBP/IsoCor) in low resolution mode (Millard et al., 2019). In addition, unlabeled chemical standards, Arabidopsis seedlings and *B. napus* leaves were additionally corrected for the contribution of naturally occurring isotopes of the carbon arising from the metabolite backbone (giving a residual molecular ^13^C enrichment of 0 for molecules labeled at natural ^13^C abundance). ^13^C-PT samples were not concerned by this additional correction since they held fully controlled ^12^C/^13^C patterns at all positions. A correction method dedicated to IsoCor was specifically created for the TMS-derivative fragment described in this study (**Table 1**), *i.e*. by considering the contribution of all atoms added by the derivatization on the isotopic measurement of the monitored fragments (chemical separation of all atoms arising from the metabolite and the derivative). All files required to run this method on IsoCor are available in the supplementary material of this article (**Supplementary files**) and all information will be updated indefinitely in the following INRAE dataverse (Dellero and Filangi, 2021). The method is publicly available and can also be run directly from GC-MS source files using the following Galaxy workflow (https://galaxy.genouest.org/, section “Metabolomics”, subsections “Conversion GCMS PostRun Analysis to IsoCor” and “Isotope Correction for mass spectrometry labeling experiments”). For this purpose, each raw GC-MS dataset should contain a column “Name” for the fragments considered and a column “Area”. The name of each fragment must be written exactly as specified in the “Metabolite.dat” (an example file is provided in the **supplementary material**). Fractional Carbon Isotopologue Distribution and mean ^13^C enrichment were directly obtained from IsoCor outputs (isotopologue_fraction and mean_enrichment) while CIDs were rescaled to Pascal’s Triangle coefficients by multiplying with 2^n^, where n is the number of carbon atoms (Millard et al., 2014). The accuracy of measurements was assessed from the closeness of experimental and predicted values for CIDs and mean ^13^C enrichments at either ^13^C-PT abundance or at natural ^13^C abundance. CID accuracy was formally defined as the mean of absolute differences between measured and predicted values for each isotopologue (Heuillet et al., 2018).

**Table 1.** Molecular ions selected for Carbon Isotopologue Distribution analysis of organic and amino acid TMS-derivatives by GC-MS.

| Compounds | TMS derivative |  | Loss |  | Molecular ion |  |  |
| --- | --- | --- | --- | --- | --- | --- | --- |
|  | Formula | mass | Formula | mass | Formula | m/z | Carbon skeleton |
| *Organic acids* |  |  |  |  |  |  |  |
| Citrate(4TMS) | C18H40O7Si4 | 480 | CH3 | 15 | C17H37O7Si4 | 465 | 1-2-3-4-5-6 |
| Citrate(4TMS) | C18H40O7Si4 | 480 | TMS-CO2, TMS-OH | 207 | C11H21O4Si2 | 273 | 1-2-3-4-5 |
| Fumarate(2TMS) | C10H20O4Si2 | 260 | CH3 | 15 | C9H17O4Si2 | 245 | 1-2-3-4 |
| Malate(3TMS) | C13H30O5Si3 | 350 | CH3 | 15 | C12H27O5Si3 | 335 | 1-2-3-4 |
| Malate(3TMS) | C13H30O5Si3 | 350 | TMS-CO2 | 117 | C9H21O3Si2 | 233 | 2-3-4 |
| Pyruvate(MEOX)(TMS) | C7H15NO3Si | 189 | CH3 | 15 | C6H12NO3Si | 174 | 1-2-3 |
| Succinate(2TMS) | C10H22O4Si2 | 262 | CH3 | 15 | C9H19O4Si2 | 247 | 1-2-3-4 |
| *Amino acids* |  |  |  |  |  |  |  |
| Asparagine(2TMS) | C10H24N2O3Si2 | 276 | TMS-CO2 | 117 | C6H15N2OSi | 159 | 2-3-4 |
| Asparagine(3TMS) | C13H32N2O3Si3 | 348 | TMS-CO2 | 117 | C9H23N2OSi2 | 231 | 2-3-4 |
| Aspartate(2TMS) | C10H23NO4Si2 | 277 | TMS-CO2 | 117 | C6H14NO2Si | 160 | 2-3-4 |
| Aspartate(3TMS) | C13H31NO4Si3 | 349 | TMS-CO2 | 117 | C9H22NO2Si2 | 232 | 2-3-4 |
| Glutamate(3TMS) | C14H33NO4Si3 | 363 | TMS-CO2 | 117 | C10H24NO2Si2 | 246 | 2-3-4-5 |
| Glutamine(3TMS) | C14H34N2O3Si3 | 362 | TMS-CO2 | 117 | C10H25N2OSi2 | 245 | 2-3-4-5 |
| Glycine(3TMS) | C11H29NO2Si3 | 291 | TMS-CO2 | 117 | C7H20NSi2 | 174 | 2 |
| Isoleucine(2TMS) | C12H29NO2Si2 | 275 | TMS-CO2 | 117 | C8H20NSi | 158 | 2-3-4-5-6 |
| Leucine(2TMS) | C12H29NO2Si2 | 275 | TMS-CO2 | 117 | C8H20NSi | 158 | 2-3-4-5-6 |
| Lysine(4TMS) | C18H46N2O2Si4 | 434 | TMS-CO2 | 117 | C14H37N2Si3 | 317 | 2-3-4-5-6 |
| Methionine(2TMS) | C11H27NO2SSi2 | 293 | TMS-CO2 | 117 | C7H18NSSi | 176 | 2-3-4-5 |
| Ornithine(4TMS) | C17H44N2O2Si4 | 420 | none |  | none |  | 1-2-3-4-5 |
| Phenylalanine(2TMS) | C15H27NO2Si2 | 309 | TMS-CO2 | 117 | C11H18NSi | 192 | 2-3-4-5-6-7-8-9 |
| Phenylalanine(2TMS) | C15H27NO2Si2 | 309 | C7H7 | 91 | C8H20NO2Si2 | 218 | 1-2 |
| Proline(2TMS) | C11H25NO2Si2 | 259 | TMS-CO2 | 117 | C7H16NSi | 142 | 2-3-4-5 |
| Serine(2TMS) | C9H23NO3Si2 | 249 | TMS-CO2 | 117 | C5H14NOSi | 132 | 2-3 |
| Serine(3TMS) | C12H31NO3Si3 | 321 | TMS-CO2 | 117 | C8H22NOSi2 | 204 | 2-3 |
| Serine(3TMS) | C12H31NO3Si3 | 321 | TMS-CH2O | 103 | C8H20NO2Si2 | 218 | 1-2 |
| Threonine(3TMS) | C13H33NO3Si3 | 335 | TMS-CO2 | 117 | C9H24NOSi2 | 218 | 2-3-4 |
| Tryptophan(2TMS) | C17H28N2O2Si2 | 348 | 2TMS-C2H2NO2 | 218 | C9H8N | 130 | 3-4-5-6-7-8-9-10-11 |
| Tryptophan(2TMS) | C17H28N2O2Si2 | 348 | C9H8N | 130 | C8H20NO2Si2 | 218 | 1-2 |
| Tryptophan(3TMS) | C20H36N2O2Si3 | 420 | 2TMS-C2H2NO2 | 218 | C12H16NSi | 202 | 3-4-5-6-7-8-9-10-11 |
| Tryptophan(3TMS) | C20H36N2O2Si3 | 420 | C12H16NSi | 202 | C8H20NO2Si2 | 218 | 1-2 |
| Tyrosine(3TMS) | C18H35NO3Si3 | 397 | C10H15OSi | 179 | C8H20NO2Si2 | 218 | 1-2 |
| Tyrosine(3TMS) | C18H35NO3Si3 | 397 | TMS-CO2 | 117 | C14H26NOSi2 | 280 | 2-3-4-5-6-7-8-9 |
| Valine(2TMS) | C11H27NO2Si2 | 261 | TMS-CO2 | 117 | C7H18NSi | 144 | 2-3-4-5 |
| Valine(2TMS) | C11H27NO2Si2 | 261 | C3H7 | 43 | C9H20NO2Si2 | 218 | 1-2 |
| α-Alanine(2TMS) | C9H23NO2Si2 | 233 | TMS-CO2 | 117 | C5H14NSi | 116 | 2-3 |
| β-Alanine(3TMS) | C12H31NO2Si3 | 305 | CH3 | 15 | C11H28NO2Si3 | 290 | 1-2-3 |

### Pathway fractional contribution

Pathway fractional contribution (FC) can be applied to a metabolic network in a metabolic and isotopic steady-state to compare the relative contribution of the considered pathway to a metabolite, as long as these pathways produced specific isotopologues for this metabolite (Buescher et al., 2015). Basically, it consists in the comparison (ratios) of mass isotopologues derived from each pathway in the final metabolite considered to determine their relative contributions (for example: isotopologues from pathway A/(isotopologues from pathway A and pathway B)). Here, FC was calculated starting from U-^13^C-pyruvate and considering malate and citrate as final metabolites. We considered two major fluxes for the biosynthesis of malate and citrate from pyruvate: the forward mode of the TCA cycle, reflecting the successive action of multiple enzyme from this cycle, and the PEPc flux, but only associated to the reassimilation of TCA cycle and PDC-derived CO_2_. Indeed, PEPc could also use ^12^CO_2_ arising independently from the decarboxylation of U-^12^C-pyruvate, leading to an unknow fraction of unlabeled malate derived from PEPc and entering the TCA cycle. To apply this methodology, we selected the timepoint 4 and 6 hours because there was a metabolic steady-state and an isotopic pseudo steady-state for isotopologue ratios. Malate biosynthesis from ^13^C_3_-pyruvate produced M1, M2 and M3 isotopologues in the fragment Malate_3TMS_C1C4, with M1 belonging to both PEPc and TCA cycle fluxes, while M2 and M3 were exclusively attributed to the TCA cycle (See result section). Interestingly, TCA cycle produced both M1 and M3 at the same time in the fragment Malate_3TMS_C1C4. This M3/(M1+M3) ratio was also observed in other fragments (Succinate_2TMS_C1C4 and Glutamate_2TMS_C2C5), relatively stable, and corresponded to the same metabolite precursors, thereby reflecting the M0 and M2 from mitochondrial active pool of acetyl-CoA (See result section). Thus, this ratio was used to calculate the proportion of M1 from Malate_3TMS_C1C4 fragment belonging exclusively to the TCA cycle based on the M3 fragment produced only by the contribution of the TCA cycle. In the following equations, *mi* refers to the isotopologue enrichment calculated with respect to the CID (sum of all *mi* for a given metabolite equal to 1). As all these calculations are relative, it is also possible to use corrected areas of mass isotopologues (outputs from IsoCor).

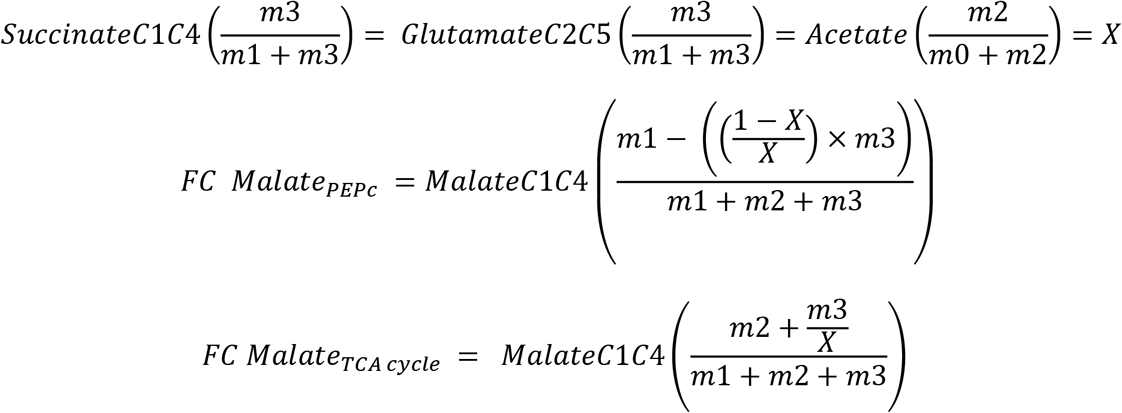

Citrate biosynthesis from U-^13^C-pyruvate produced M1, M2, M3 and M4 isotopologues in the fragment Citrate_4TMS_C1C6, with M1 belonging exclusively to PEPc, M3 belonging to both PEPc and TCA cycle and M2 and M4 belonging exclusively to TCA cycle (See result section). Therefore, FC were calculated as follow:

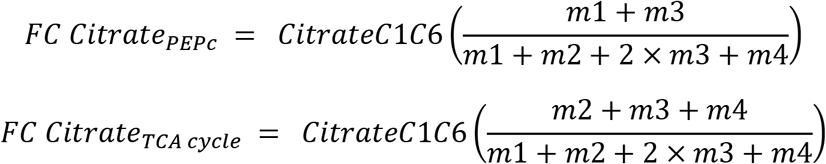

Finally, the FC of TCA cycle and PDC-derived CO_2_ to photosynthesis was calculated using serine fragments and the ratio “*X”* as a proxy for the ^13^C/^12^C ratio of this CO_2_ fraction. We did not consider the contribution of plastidial PDC here since a recent ^13^C-MFA of tobacco leaves showed that this flux was relatively negligible compared to the mitochondrial PDC flux (Chu et al., 2022). Considering the isotopic and metabolic steady-state for serine fragments, the mean ^13^C enrichment of *X* should be observed in serine if the mitochondrial fraction of CO_2_ accounted for 100% of the CO_2_ fixed by photosynthesis. Thus, FC was calculated as follow:

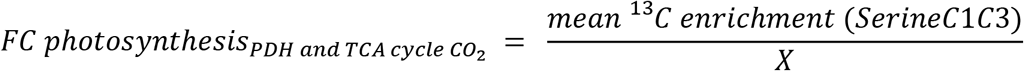

For this, mean ^13^C enrichment of serine was recalculated from M1 isotopologue of either Serine_2TMS_C1C2 or Serine_2TMS_C2C3, since serine was only labeled in the C2 of these fragments based on the hypothesis validated by our results.

### Statistical analysis

Statistical comparisons were performed using either confidence intervals (comparison of a group of value against one reference value), a Student-test (comparison of 2 groups of values) or an ANOVA followed by a post-hoc Tukey-HSD test (comparison of 3 or more groups of values) with a level of p-value <0.05 by using the publicly available Rstudio software and dedicated packages (RStudio Team, 2016).

## 3 RESULTS

### Evaluation of CID measurements by GC-MS

The proposed workflow (**Figure 1**) consisted in three steps: 1) the build-up of a method to identify and select abundant molecular ions deriving from TMS-derivatives with a large number of carbons; 2) the production of two reference ^13^C-labeled samples with a controlled pattern for CID (^13^C-PT) using *E. coli* as a metabolic factory; 3) the validation of TMS-derivatives mass fragments for CID measurements with our GC-MS method for applications in plants using ^13^C-PT reference samples and unlabeled and ^13^C-labeled plant samples. First, we combined a literature survey analyzing common pathway of TMS-derivative mass fragmentations with the injection of chemical standards in our analytical system (Lai and Fiehn, 2018;Okahashi et al., 2019;Harvey and Vouros, 2020). We selected the most abundant fragments maximizing the information on carbon backbones (**Table 1**). Their chemical identity was further confirmed by the observed CID pattern with our two reference ^13^C-PT samples. Next, we produced reference ^13^C-labeled samples with *E. coli* (^13^C-PT samples) using an accurate equimolar mixture of all ^13^C isotopomers of acetate as previously described (Millard et al., 2014). This strategy produced metabolites with fully controlled and predictable ^13^C-labeling patterns, *i.e*. a mean ^13^C enrichment close of 50% (49.9 % ± 0.8 in our study based on NMR analysis) and a CID following coefficients from Pascal’s triangle (binomial distribution) (Millard et al., 2014). From *E. coli* cell harvest, two specific ^13^C-PT samples were produced during the extraction: i) a ^13^C-PT full metabolite extract containing all polar metabolites for the evaluation of CID for organic acids; ii) a ^13^C-PT protein hydrolysates extract containing proteogenic amino acids in high concentrations for the evaluation of CID for amino acids. While this strategy allowed to maximize the amount of soluble proteogenic amino acids detectable, it prevented the evaluation of glutamine and asparagine due to the hydrolysis process (Kambhampati et al., 2019).To facilitate comparison of CIDs between the different amino acids, we rescaled the measured ^13^C-enrichments among isotopologues of each fragment into binomial distribution, thus reflecting the average number of isotopomers constituting each isotopologue (**Figure 2** and Material and Methods). In the following figures, GC-MS fragments will be denoted according to the considered metabolite, its TMS-derivative analyzed and the metabolite carbon backbone (for example, Glutamate_3TMS_C2C5 means that the fragment contains the carbon C2-C3-C4-C5 (**Table1**)).

**Figure 2.**
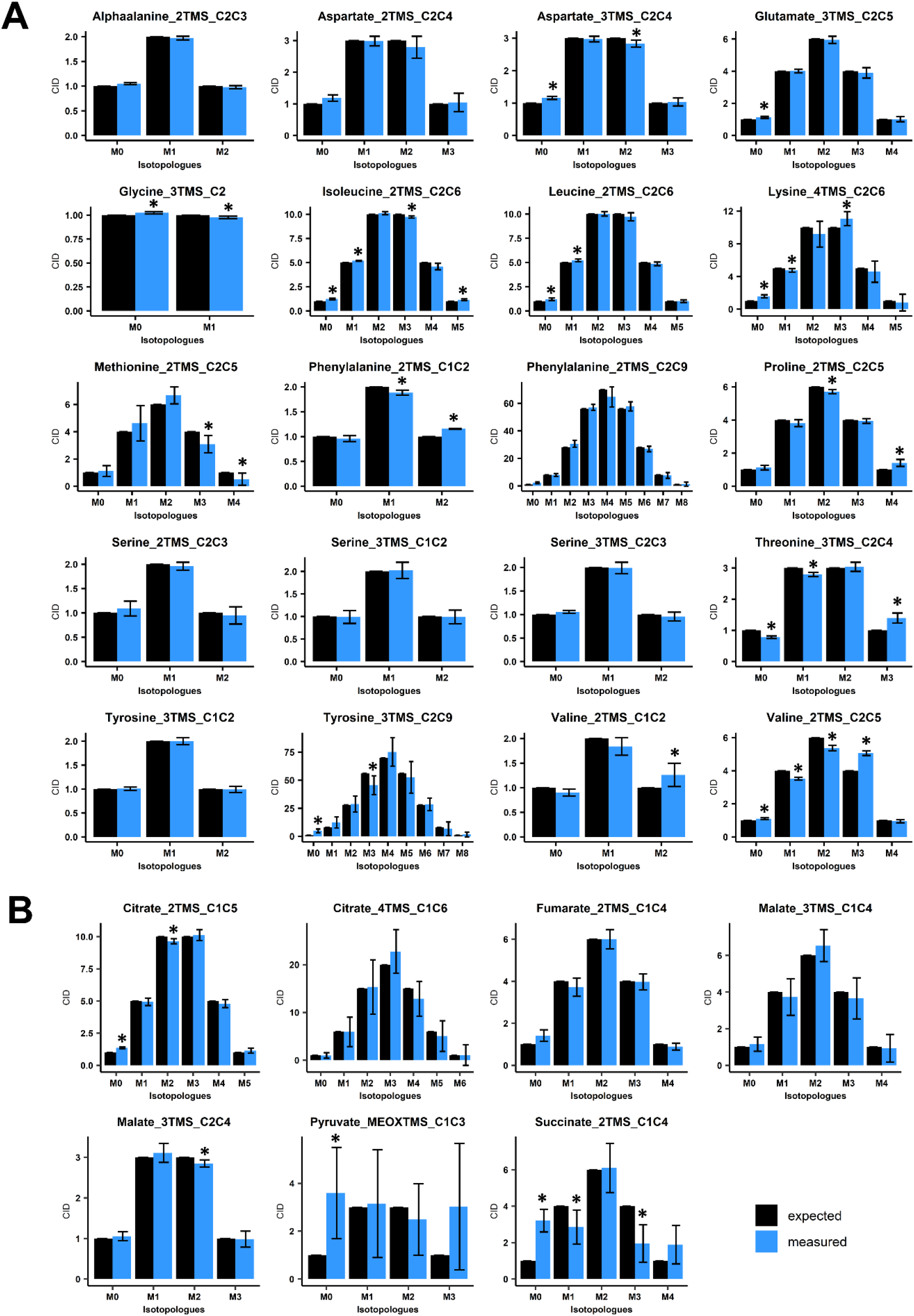
Evaluation of Carbon Isotopologue Distribution (CID) measurements for organic and amino acids by GC-MS using ^13^C-PT reference samples. **A**, Evaluation of amino acids using ^13^C-PT protein hydrolysates. **B**, Evaluation of organic acids using ^13^C-PT full metabolites extracts. Black bars correspond to expected CIDs and blue bars to the measured CIDs. The results are presented as the mean ± SD of four independent biological replicates. Statistical differences between predicted and measured CID for each isotopologue of each fragment are denoted with asterisks (*) and were established by considering the 95% confidence intervals of the measured CID. GC-MS fragments are denoted according to the considered metabolite, its MEOX/TMS-derivatives analyzed and the metabolite carbon backbone (for example, Glutamate_3TMS_C2C5 means that the fragment comes from glutamate(3TMS) and contains the carbon C2-C3-C4-C5 (**Table 1**)).

Overall, all the considered amino acids were detected in the ^13^C-PT protein hydrolysates except tryptophan (**Figures 2A, S1A**). We found very a high level of correlation between the measured CID (blue bars) and the expected CID (black bars) for most of the amino acid fragments considered in our method. CID measurements for Methionine_2TMS_C2C5, Lysine_4TMS_C2C6 and Tyrosine_2TMS_C2C9 showed a high variability, leading to small misestimations for some isotopologues (**Figure 2A**). For methionine, this could be a consequence of its very low abundance in the ^13^C-PT protein hydrolysates compared to the others amino acids (**Figure S1A**). The most important and significant errors for CID measurements of amino acids concerned Proline_2TMS_C2C5 (overestimation of M4), Threonine_3TMS_C2C4 (overestimation of M3 at the expense of M0 and M1), Valine_2TMS_C1C2 (overestimation of M2) and Valine_2TMS_C2C5 (overestimation of M3 at the expense of M1 and M2). Considering the recent establishment of a ^13^C-positional isotopomer approach for glutamate to analyze its C1 carbon in connection with PEPc activity (Lima et al., 2021), we also analyzed the CID for the two low abundant fragments Glutamate_3TMS_C1 (CO_2_-TMS, M0 at m/z 117) and Glutamate_3TMS_C1C5 fragment (CH3 loss, M0 at m/z 348) (**Figure S1B**). However, they both showed large and significant misestimations of their CIDs, thus questioning the accuracy of this isotopomer-based method. The analysis of organic acids from ^13^C-PT full metabolites extract showed relatively good CID for Citrate_2TMS_C1C5, Malate_3TMS_C2C4 and Fumarate_2TMS_C1C4 (**Figure 2B**). Citrate_4TMS_C1C6 and Malate_3TMS_C1C4 showed also relatively good CID but they had a higher variability compared to their respective C1C5 and C2C4 fragments. There was a very high overestimation of M0 in Succinate_2TMS_C1C4, a contamination already reported for this ^13^C-PT full metabolites extract (Millard et al., 2014), which prevented its validation by the workflow. For Pyruvate_MEOXTMS_C1C3, the variability was too high to assess the CID measurements, perhaps due to its weak derivatization yield and low abundance in the^13^C-PT full metabolites extract (**Figure S1A**). To further assess the distribution of measurement errors among isotopologues, we analyzed the CID accuracy, using threshold values of ±0.05 (**Figure 3A**). Overall, the bias of measurement for each isotopologue of each metabolite was relatively low except for methionine, valine, threonine, succinate and pyruvate. Interestingly, these very high levels of accuracy for CID measurements were converted into very high levels of accuracy for fractional mean ^13^C enrichment measurements, leading to very small shifts between expected (0.499 ±0.08) and measured values (below 1% of error), except again for methionine, valine, threonine, succinate and pyruvate (**Figure 3B**). Nevertheless, Malate_3TMS_C1C4, Lysine_2TMS_C2C6 and Serine_2TMS_C2C3 showed important variability for mean ^13^C enrichment measurements although no strong biases in their CID measurements were previously observed. Conversely, fractional mean ^13^C enrichment of Phenylalanine_2TMS_C1C2 and Proline_2TMS_C2C5 were slightly overestimated and reflected the overestimation of their respective isotopologues M2 and M4.

**Figure 3.**
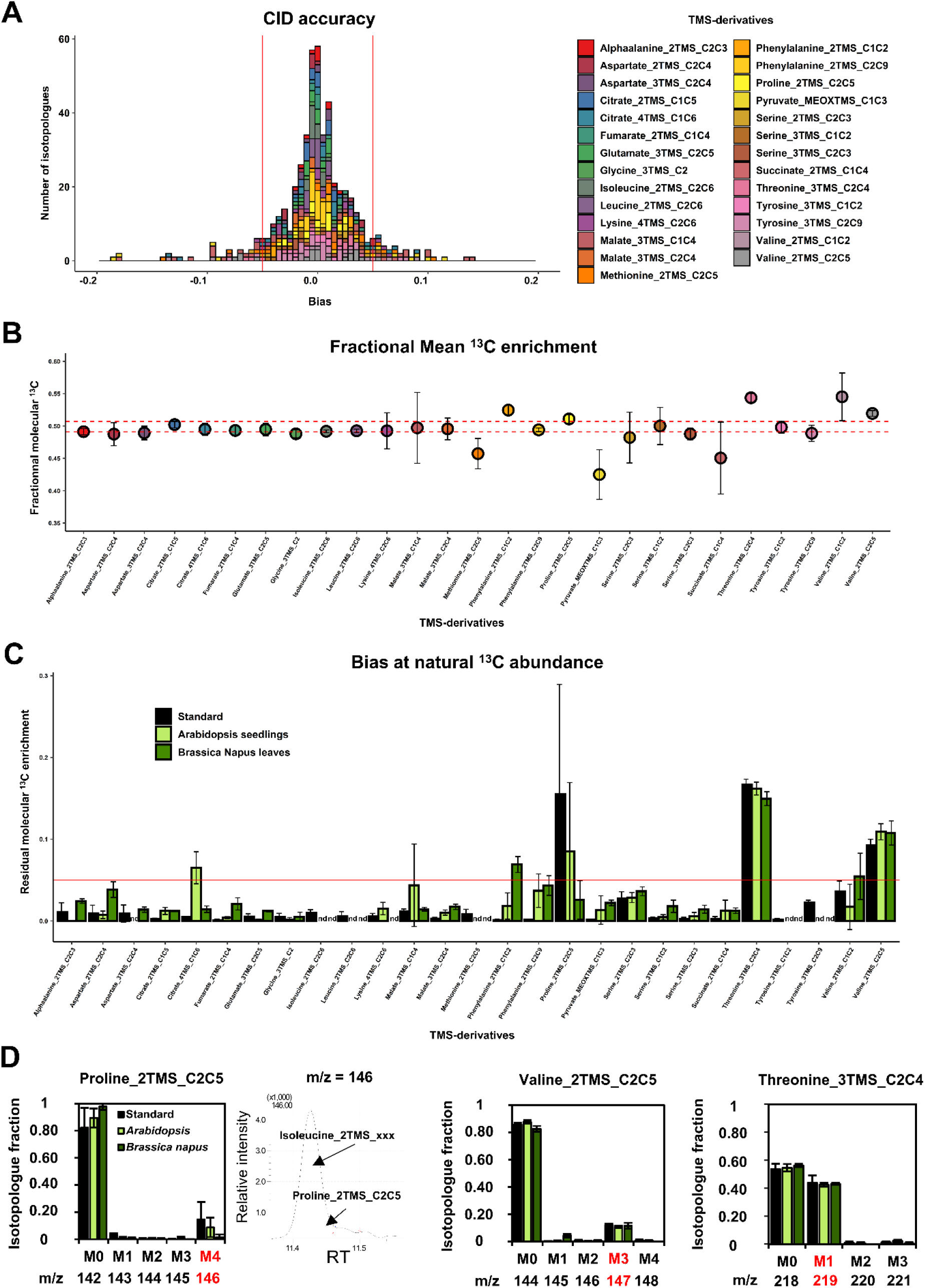
Evaluation of the measurement accuracy for CID and mean ^13^C enrichment at ^13^C-PT and ^13^C natural abundance in plants. **A**, CID accuracy and **B**, Mean ^13^C enrichment for the selected TMS-derivatives of organic and amino acids using ^13^C-PT samples. **C**, Residual mean ^13^C enrichment measured in chemical standards, Arabidopsis seedlings and *B. napus* leaves after correction for isotope natural abundance. **D**, Isotopologue-specific biases identified at ^13^C natural abundance for proline, valine and threonine fragments. The results are presented as the mean ± SD of four independent biological replicates or independent preparations for commercial chemical standards.

Then, we evaluated the suitability of our GC-MS method for application in plants. First, we analyzed measurement accuracy at natural ^13^C abundance using either seedlings from Arabidopsis or leaves from *B. napus*. To do so, we measured the fractional residual molecular ^13^C enrichment, *i.e* the fractional ^13^C enrichment remaining after removing the contribution of natural abundance for all isotopes (a value of 0 is expected if no contamination occurs) (**Figure 3C**). Among the metabolites detected in the plant samples, most of the fragments for organic and amino acids showed relatively high accuracies: mean values ranged between and 0.005 and 0.03 (**Figure 3C**). Some discrepancies were observed between Arabidopsis seedlings and *B. napus* leaves for Citrate_4TMS_C1C6 and Malate_3TMS_C1C4 (higher accuracy for *B. napus*), and for AlphaAlanine_2TMS_C2C3, Aspartate_2TMS_C2C4, Fumarate_2TMS_C1C4 and Serine_3TMS fragments (higher accuracy for Arabidopsis). Residual molecular ^13^C enrichments for Phenylalanine_2TMS fragments were close to 0.05 but only in plants samples. The major errors at natural ^13^C abundance concerned Proline_2TMS_C2C5, Threonine_3TMS_C2C4 and Valine_2TMS_C2C5 and were observed in both chemical standards and unlabeled plant extracts, confirming the occurrence of systematic matrix-independent analytical bias. A detailed survey showed that the isotopic cluster of Proline_2TMS_C2C5 was contaminated in the M4 isotopologue (m/z 146) by a mass fragment deriving from near elution of Isoleucine_2TMS, and correlated with the M4 overestimation previously observed in ^13^C-PT protein hydrolysate sample (**Figures 2, 3D**). Similarly, the isotopic cluster of Valine_2TMS_C2C5 was contaminated in the M3 isotopologue (m/z 147) while Threonine_3TMS_C2C4 showed a surprising very high abundance of the M1 isotopologue (40% of the CID).

### U-^13^C-pyruvate incorporations into leaf discs and qualification of metabolic and isotopic states

We used our validated MS method to explore the light/dark regulation of TCA cycle and PEPc fluxes in *B. napus* leaves, with a special emphasize on day/night citrate metabolism and the cyclic/non-cyclic modes of the TCA cycle. For this purpose, we selected U-^13^C-pyruvate as a metabolic probe since the mitochondrial Pyruvate Dehydrogenase complex (PDC) will be able to produce simultaneously ^13^C_2_-acetyl-CoA molecules suitable for the investigation of the TCA cycle and ^13^CO_2_ molecules suitable for the investigation of PEPc activity through the combined action of carbonic anhydrase (H^13^CO_3_^-^).

Following transient kinetics of 10 mM U-^13^C-pyruvate incorporation in either light or dark conditions for up to 6 hours, the analysis of mean ^13^C enrichment for all conditions revealed a significant increase for fragments of alphaalanine, aspartate, citrate, glutamate, glycine, malate, pyruvate, serine and succinate along time (**Figure 4A**, **Table S2**; **Table S4** for statistical analysis). There was a very high biological variability and small ^13^C enrichment for fumarate fragment. Overall, most of these enrichment dynamics remained relatively equivalent between light and dark conditions for all the fragments considered, except for glycine and serine, which were almost exclusively enriched in light conditions. These results were surprising since the commitment of an unlabeled stored ^12^C-citrate during light conditions was expected to introduce an isotopic dilution compared to dark conditions, that could have been reflected in citrate, glutamate and succinate notably. Considering glycine and serine, the importance of the light-dependent photorespiratory pathway for their production in leaves clearly suggested that a significant part of ^13^CO_2_ released was reassimilated by photosynthesis. A statistical analysis revealed that Glycine_3TMS_C2, Serine_3TMS_C1C2 and Serine_3TMS_C2C3 reached an isotopic pseudo steady-state starting after 2 hours of illumination (**Table S4**). Other fragments showed a relatively linear evolution of their mean ^13^C enrichment, except for Pyruvate_MEOXTMS_C1C3 which rapidly reached an isotopic steady-state after 0.5 hours only. Interestingly, labeling data from Pyruvate_MEOXTMS_C1C3 and Succinate_2TMS_C1C4 fragments had a very good resolution (weak analytical and biological variability) compared to previous results obtained with ^13^C-PT full metabolites extract.

**Figure 4.**
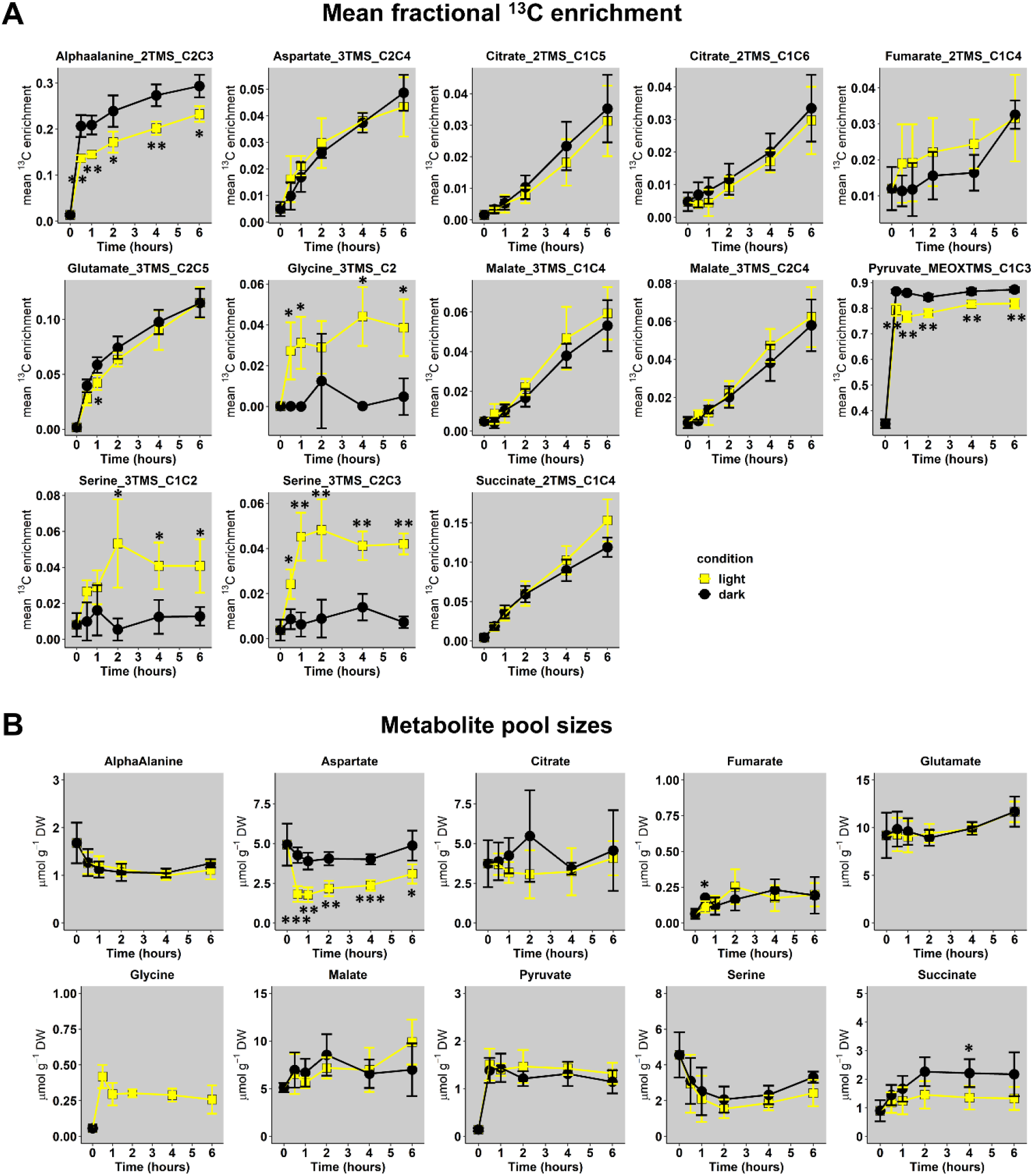
Dynamics of U-^13^C-pyruvate incorporation in *Brassica napus* leaf discs exposed to light or dark conditions. **A**, Mean fractional ^13^C enrichment and **B**, Metabolite pool sizes. Mean fractional ^13^C enrichment were monitored with GC-MS, while absolute quantification of organic and amino acids was performed respectively with GC-FID and HPLC-UV. The results are presented as the mean ± SD of four independent biological replicates. Statistical differences between dark and light conditions are denoted with asterisks according to a Student test (*, p-value <0.05; **, p-value <0.01; ***, p-value <0.001). The complete dataset is available in **Table S2, S3**. Supplementary statistics (ANOVA and post-hoc Tukey-HSD tests) are available in **Table S4, S5**.

Next, we performed quantification of total pool sizes for these metabolites to give an approximate indication whether the leaf metabolism was in a metabolic pseudo steady-state (plant cells are extensively compartmented) (**Figure 4B, Table S3**). This was done by GC-FID for organic acids and UPLC-UV for amino acids using AccQTag methodology. Indeed, the response of TMS-derivatives for amino acids is too unstable along a GC-MS run (**Figure S3**) and absolute quantification of metabolites using their CID masses have to be achieved according to internal isotopic standards for each metabolite (not our case in the present study) (Heuillet et al., 2020;Evers et al., 2021). Overall, total metabolite pool sizes were relatively different between each other’s for any given time but there was no significant difference between light and dark conditions, except for Aspartate. Glycine was only quantified in light conditions (not detected in “dark condition” samples with UPLC-UV). Interestingly, a statistical analysis showed that all the total metabolite pool sizes reached a significant pseudo steady-state starting after 1 hour (values belonging to similar statistical groups in **Table S5**). To summarize, we found some similar ^13^C enrichment dynamics for TCA cycle derived metabolites in both light and dark conditions combined to similar total metabolite pool sizes at a metabolic pseudo steady-state.

### Metabolic network considerations and dynamics of CIDs for TCA cycle-derived metabolites

According to our ^13^C-labeling strategy and the metabolites detected with our MS method, we constructed a metabolic network containing all the possible isotopomers that will be produced from two “turns” of the TCA cycle and the PEPc activity, including the reassimilation of TCA cycle and PDC-derived CO_2_ (**Figure 5A**). In this network, the second “turn” reflected the commitment of a molecule that has already experienced all reactions of the TCA cycle to another round of reactions (Orange and blue labels to compare acetyl-CoA-derived labeling from 1^st^ and 2^nd^ “turn”). Based on previous works (Abadie and Tcherkez, 2019), the reverse reaction of the fumarase was also considered in the network, although it had no effect on the M1 isotopologue monitored by mass spectrometry (C4 and C1 isotopomers are both detected as M1 isotopologues in our study). The involvement of a light-dependent stored ^12^C-citrate pool was not represented here since it only provided M0 isotopologues (no additional isotopologues to explain). The activity of the malic enzyme, converting malate to pyruvate, was also not considered here since it may have been inhibited due to the high incorporation of pyruvate. Overall, the PEPc-dependent reassimilation of TCA cycle and PDC-derived CO_2_ was expected to introduce M1 isotopologues for Malate_3TMS_C1C4 and M1 and M3 isotopologues for Citrate_4TMS_C1C6. Conversely, the non-cyclic mode of the TCA cycle should introduce M2 isotopologues to Citrate_4TMS_C1C6, Glutamate_2TMS_C2C5 and Succinate_2TMS_C1C4 (incorporation of M2 acetyl-CoA arising from U-^13^C-pyruvate) while the cyclic mode should introduce M2 isotopologue to malate, followed by M1, M3 and M4 isotopologues for all these metabolites if the M2 isotopologues were committed again to the TCA cycle (second “turn”).

**Figure 5.**
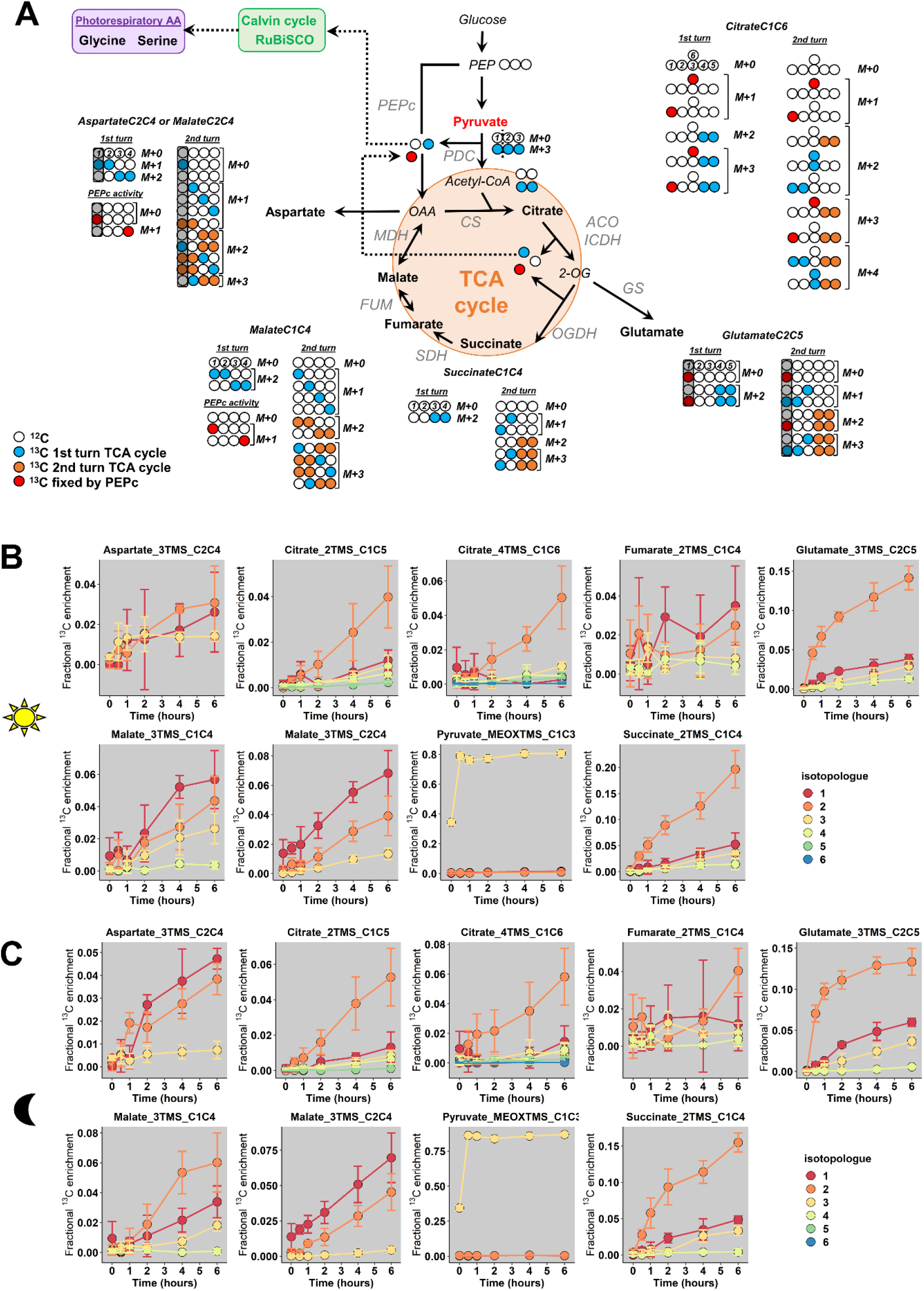
Fractional ^13^C enrichment at the isotopologue level for TCA-cycle derived metabolites. **A**, Metabolic network considered based on the ^13^C-labeling strategy. **B**, Light conditions and **C**, Dark conditions. The M0 isotopologue was not represented in the graphs to facilitate the visualization of low-enriched isotopologues. The results are presented as the mean ± SD of four independent biological replicates. The complete dataset is available in **Table S2**. In the proposed network, the second “turn” reflected the commitment of a molecule that has already experienced all reactions of the TCA cycle to another round of reactions (Orange and blue labels to compare acetyl-CoA-derived labeling from 1^st^ and 2^nd^ “turn”). Note that M0 istopologues of all metabolites will still continue to produce M0 istopologues at an important rate after a “turn” of the TCA cycle, given the ^13^C-enrichment for acetyl-CoA active pool. PEPc, Phosphoenolpyruvate carboxylase; PDC, Pyruvate dehydrogenase complex, CS, Citrate synthase; ACO, Aconitase; ICDH, Isocitrate dehydrogenase; OGDH, 2-oxoglutarate dehydrogenase; SDH, Succinate dehydrogenase complex; FUM, Fumarase; MDH, Malate dehydrogenase.

Analysis of the dynamic of CIDs showed that mean ^13^C enrichment of citrate, glutamate and succinate was essentially driven by an increase of the M2 isotopologue for light and dark conditions, thus reflecting the conventional non-cyclic mode of the TCA cycle in both conditions (**Figure 5B,C**). However, there was also an increase of M1 and M3 isotopologues for Glutamate_2TMS_C2C5 and Succinate_2TMS_C1C4 starting after 2 hours while M1, M3 and M4 isotopologues of Citrate_4TMS_C1C6 and Citrate_2TMS_C1C5 were increased starting from 4 hours (**Figure 5B,C**). Therefore, the cyclic mode of the TCA cycle was necessarily operating during both light and dark conditions. Nevertheless, M1 and M4 of Citrate_4TMS_C1C6 were essentially increased in dark conditions compared to light conditions after 4 hours, perhaps reflecting a higher turn-over for the TCA cycle in dark conditions. There was a very high variability for Fumarate_2TMS_C2C4 CID in both conditions, which prevented its use for a reliable isotopic investigation. Interestingly, Malate_3TMS_C1C4 fragment showed a higher enrichment of M1 compared to M2 in the light condition while M2 remains higher than M1 in the dark condition. This shift may reflect an increase of the PEPc activity under light condition. Since there was no difference of mean ^13^C enrichment between light and dark conditions for this fragment (**Figure 4A**), this result reinforced the importance of analyzing specifically CIDs for the interpretation of labeling data from TCA cycle. In addition, the CID of Malate_3TMS_C2C4 showed a similar dynamic for both light and dark conditions, hence ruling out the use of this fragment to disentangle PEPc and TCA cycle contribution to malate production (**Figure 5B,C**). Aspartate_3TMS_C2C4 showed a very high variability for the light condition, but the CID dynamic in the dark condition was similar to the CID dynamic of Malate_3TMS_C2C4. Thus, both fragments seemed to mimic oxaloacetate labeling dynamics. Pyruvate_MEOXTMS_C1C3 was only enriched with the M3 isotopologue (nearly 80% M3/20% M0), thereby confirming a negligible activity of the malic enzyme in our metabolic network.

### Pathway fractional contribution for TCA cycle and PEPc-dependent reassimilation of TCA cycle and PDC-derived CO_2_ to either malate and citrate biosynthesis

We took advantage of the pathway-specific mass isotopologues produced respectively by the two “turns” of the TCA cycle and PEPc to calculate pathway fractional contributions (FC) for pyruvate-to-malate and pyruvate-to-citrate (**Figure 6A**). As PEPc activity could also be driven by other sources of CO_2_ (photorespiration and ambient air) diluting the final ^13^C/^12^C ratio for HCO_3_^-^, our results presented here only reflected the PEPc activity devoted to the reassimilation of TCA cycle and PDC-derived ^13^CO_2_. We used only the timepoints 4 and 6 hours, because all the fragments considered for these FC were mostly visible at these moments. In addition, pathway FC can only be applied if the metabolic network is at both metabolic and isotopic pseudo steady-states. Here, we observed a rapid metabolic pseudo steady-state (**Figure 4B**, **Table S5**) and a relatively negligible evolution of the ratios for the considered isotopologues between 4 and 6 hours (similar evolutions of M1, M2 and M3 isotopologues for malate between 4 and 6 hours for example). Thus, the relative contribution of PEPc and TCA cycle to either malate and citrate remained in the same order of magnitude between 4 and 6 hours (**Figure 5B,C**). An important issue to address for calculating FC concerned the M1 from Malate_3TMS_C1C4, which monitored at the same time PEPc and TCA cycle dependent isotopomers (**Figure 5A**). To deal with, we took advantage of Succinate_2TMS_C1C4 and Glutamate_2TMS_C2C5 fragments which both harbored M1 and M3 isotopologues that are the precursors of TCA cycle-dependent M1 and M3 of Malate_3TMS_C1C4 (**Figures 5A and 6A**). Analysis of the M3/(M1+M3) ratio for these two fragments showed a remarkable stability between dark and light conditions, thus reflecting a stable M2 enrichment of 40% for the active pool of acetyl-CoA (**Figure 6B** and Equations). This result also confirmed that the isotopologue ratios considered here were at an isotopic pseudo steady-state. Using this ratio, we calculated the FC of PEPc and TCA cycle for pyruvate-to-malate and pyruvate-to-citrate respectively (**See equations**). We found that TCA cycle had a significant four-fold higher contribution for pyruvate-to-malate or pyruvate-to-citrate compared to PEPc in all conditions (around 80% versus 20%). Thus, PEPc-dependent reassimilation of TCA cycle and PDC-derived CO_2_ was relatively low compared to the forward cyclic mode of the TCA cycle in *B. napus* leaf discs under either light or dark conditions. Interestingly, there was a 1.6-fold increase of the FC of PEPc for pyruvate-to-malate from dark to light conditions (14% versus 23%), although it remained not statistically significant (high variability, t-test p-value =0.08). Conversely, PEPc contribution for pyruvate-to-citrate was significantly 1.5-fold higher in dark condition than in light condition (26% versus 17%). These last results suggested a possible light/dark regulation of PEPc activity and a higher commitment of PEPc-derived carbons to the forward part of the TCA cycle in the dark.

**Figure 6.**
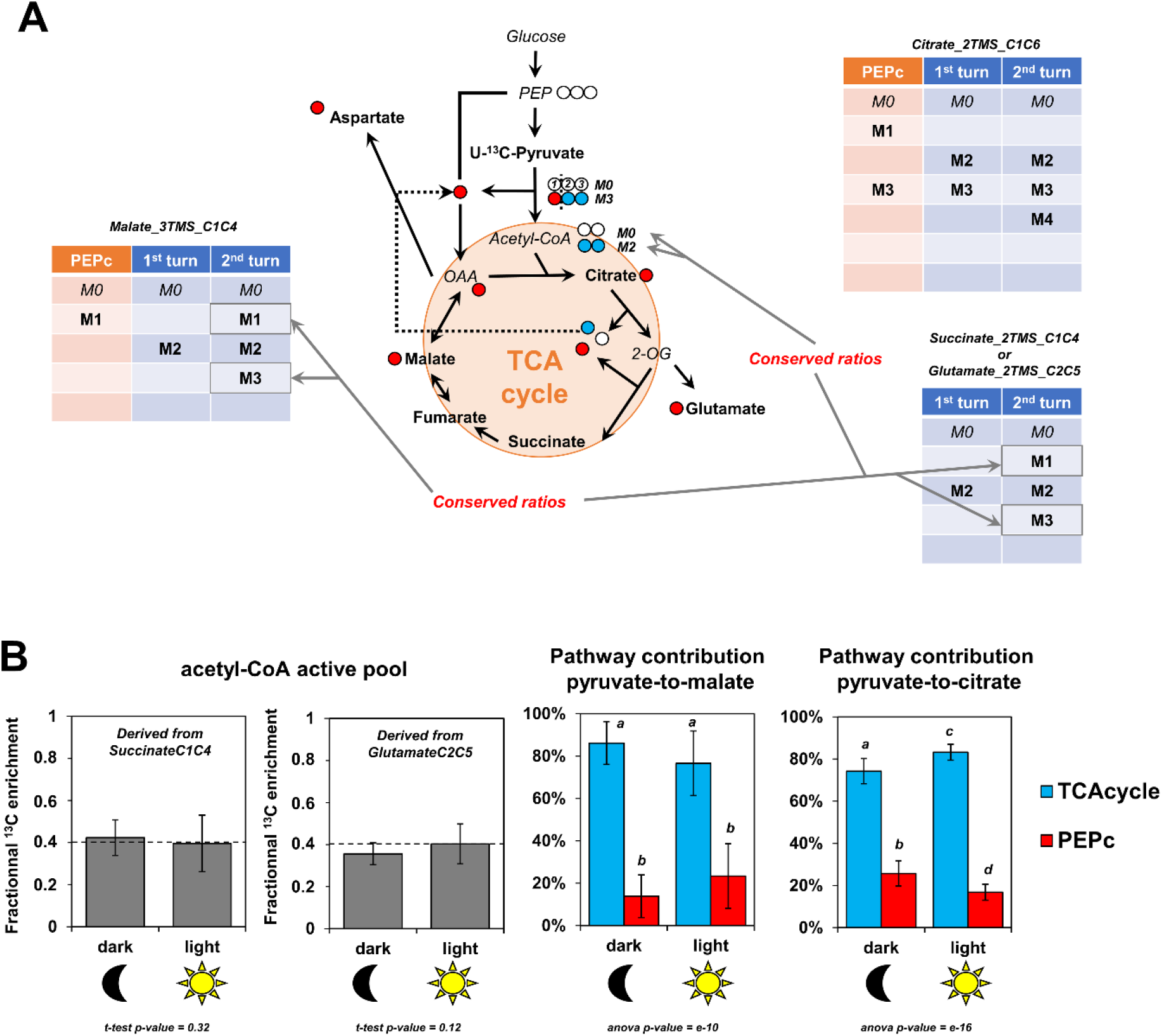
Fractional contribution of TCA cycle and PEPc-dependent reassimilation of mitochondrial CO_2_ for pyruvate to malate and pyruvate to citrate in light and dark conditions. **A**, Pathway-specific mass isotopologues considered for the calculations. **B**, Fractional contribution of PEPc and TCA cycle to malate and citrate based on pathway-specific mass isotopologues/isotopomers. For these calculations, only the timepoint T4 and T6 hours were used and pooled (metabolic steady-state and isotopic steady-state for isotopologue ratios). The results are presented as the mean ± SD of four independent biological replicates. Statistical comparison of acetyl-CoA enrichment for light and dark conditions was performed with a Student-test, while fractional contributions were compared with an ANOVA followed by a post-hoc Tukey-HSD test (different letters indicate groups that were separated with a p-value <0.05).

### Contribution of TCA cycle and PDC-derived CO_2_ to photosynthesis

To finish, we explored the isotopologue distribution of glycine and serine, some amino acids which were previously identified as a proxy for photosynthetic reassimilation of PDC and TCA cycle-derived ^13^CO_2_ (**Figure 7A**). We did not consider the contribution of plastidial PDC here since a recent ^13^C-MFA of tobacco leaves showed that this flux was relatively negligible compared to the mitochondrial PDC flux (Chu et al., 2022). Our results showed that the increase of mean ^13^C enrichment for the three considered fragments in light condition came from an increase of their M1 isotopologue. Given that the light-dependent photorespiratory enzymes glycine decarboxylase and serine:hydroxymethyl aminotransferase convert 2 glycine into 1 serine, we wanted to know whether the fragments Serine_2TMS_C1C2 and Serine_2TMS_C2C3 were labeled in the same or different carbons, thus revealing if the C1 of glycine was also labeled (**Figure 7B**). By reviewing all the possible scenarios with different combinations of labeled glycine, we showed that the CID associated to our fragments (isotopologue occurrence) could only be produced with one molecule of glycine labeled at the C2 and an unlabeled molecule of glycine, leading to the production of a molecule of serine labeled only at the C2 and an unlabeled CO_2_ molecule. Since we have shown that glycine and serine were both at the isotopic and metabolic steady-state between 2 and 6 hours (**Figure 4**, **Table S4**, **Table S5**), we used serine fragments to evaluate the contribution of mitochondrial decarboxylations to the photosynthetic activity. For this purpose, the steady-state enrichment of acetyl-CoA calculated before was used as a proxy of the ^13^C/^12^C ratio of the TCA cycle and PDC-derived CO_2_ (**See equations**) given its preponderant contribution for light respiration (CO_2_ emissions) (Tcherkez et al., 2005). After recalculating the mean ^13^C-enrichment of serine from both serine fragments (C1-C2 or C2-C3), we showed that the light-dependent assimilation of TCA cycle and PDC-derived decarboxylations could represent around 5% of net photosynthesis (**Figure 7C**).

**Figure 7.**
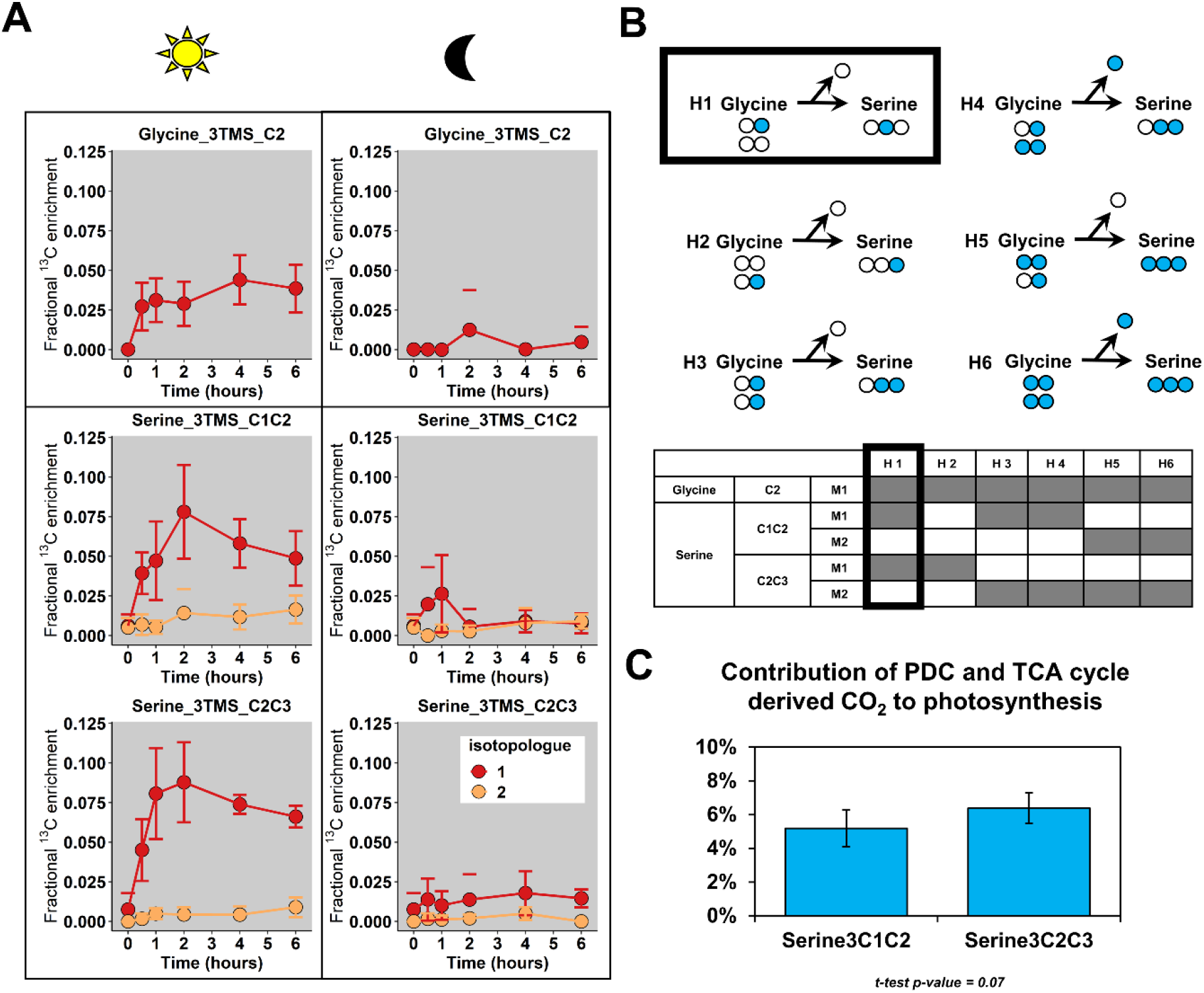
Fractional contribution of PDC and TCA-cycle derived decarboxylations to photosynthesis. **A**, Fractional ^13^C enrichment at the isotopologue level for glycine and serine. **B**, Working hypotheses to explain the photorespiratory-dependent labeling pattern of glycine and serine. **C**, Estimation of the fractional contribution of PDC (pyruvate dehydrogenase complex) and TCA-cycle CO_2_ to photosynthesis with Serine (expressed in %). For these calculations, only the timepoint T4 and T6 hours were used and pooled (metabolic and isotopic steady-state, assumption with ^13^C/^12^C of CO_2_ released). The results are presented as the mean ± SD of four independent biological replicates. Statistical comparison of the two serine fragments was performed with a Student-test.

## 4 DISCUSSION

The estimation of metabolic fluxes in photosynthetic organisms represents an important challenge that has gained interest over the last decade with the development of ^13^C-INST-MFA. This approach requires a high level of accuracy for CID measurements to avoid propagation of small measurement errors to large errors in estimated fluxes (Antoniewicz et al., 2006). Considering the need to perform high throughput measurements at isotopically non-stationary steady-state, MS has been preferentially selected by the field in the recent years to measure CID in organic and amino acids. However, the suitability of TMS-derivative molecular ions for these metabolites has never been evaluated up to the isotopologue level, *i.e*. by monitoring the measurement accuracy for each possible mass isotopologue and its impact on the final CID. The use of a validated MS methodology will help to routinely investigate ^13^C-labeling dynamics within plant TCA cycle and strengthen our knowledge about its functioning.

### A robust methodology to evaluate CID measurements and identify analytical and matrix biases

In this study, we proposed a workflow to assess the measurement accuracy of CID from TMS-derivatives of organic and amino acids by GC-MS for applications in plants, based on tailor-made *E. coli* ^13^C biological standards with controlled patterns of CID (^13^C-PT) and plant samples at natural ^13^C-abundance (**Figure 1**). Overall, most of the TMS-derivatives selected (Table 1) were validated, harboring remarkable accuracies leading to very small biases in the measurement of CIDs (below 0.05) and fractional mean ^13^C enrichments (below 0.01). Nevertheless, we identified three important biases at the isotopologue level. The first bias concerned the contamination of the isotopic cluster of Proline_2TMS_C2C5 at the m/z 146 (**Figure 3D**), which was previously reported with chemical standards (Antoniewicz et al., 2006). Here, we showed that the contamination at m/z 146 could occur from the co-elution with isoleucine: separation by 0.02 s in our chromatograms, *i.e* 0.33 °C along the temperature gradient (+10°C.min^−1^). Interestingly, this contamination had a weak impact on the mean ^13^C enrichment measured (**Figure 3B**), suggesting that this fragment may still be used, as long as the ^13^C-labeling strategy can prevent the production of M4 isotopologues. The second bias identified concerned the contamination of the isotopic cluster of Valine_2TMS_C2C5 at the m/z 147. During the fragmentation process, a siliconium ion (loss of a methyl in a TMS group) can operate a cyclization by attacking another TMS group of the same molecule, thereby producing an ion [(CH^3^)^3^Si-O-Si(CH^3^)^2^]^+^ of the same mass 147 (Harvey and Vouros, 2020). We also detected a significant abundance of an m/z 246 (siliconium ion) in the fragmentation spectra of Valine_2TMS, which seems to support this hypothesis for the contamination (**Figure S2**). The third bias concerned the very high level of M1 isotopologue detected for Threonine_3TMS_C2C4. In its fragmentation spectra, we also found a M1 isotopologue contamination for the peak m/z 291, an important fragment for the identification of Threonine_3TMS by GC-MS (**Figure S2**, NIST database). Thus, this contamination could be due to: i) a co-elution of a contaminant chemically similar to threonine; ii) a strong analytical isotopic effect. The use of GC coupled to high-resolution MS may help to resolve this analytical issue. Besides this, we could not evaluate Succinate_2TMS_C1C4 with our workflow, due to a previously reported *E. coli*-dependent contamination of ^13^C-PT samples (Millard et al., 2014). However, this fragment showed small biological and analytical variabilities in our ^13^C-labeling experiments and provided M3/(M1+M3) ratios similar to those calculated with Glutamate_2TMS_C2C5 (**Figure 6B**), hence partially validating its use for ^13^C-MFA in plants. Using our validated MS method, we provided relevant information about the activity of the TCA cycle and citrate metabolism in *B. napus* leaves that counterbalanced some conclusions from previous works (Tcherkez et al., 2009;Gauthier et al., 2010;Sweetlove et al., 2010;Tcherkez et al., 2017;da Fonseca-Pereira et al., 2021).

### Does a stored citrate pool really contribute to the mitochondrial TCA cycle activity in *Brassica napus* leaves in a light/dark-dependent manner ?

The hypothesis about the role of night-stored citrate molecules was initially proposed based on ^13^CO_2_ labeling incorporation in detached leaves of *Xanthium stumarium* in order to fit a model based on “transmission coefficients” (Tcherkez et al., 2009). A second work combined ^13^CO_2_ with ^5^NH_4_ ^15^NO_3_ labeling experiments in *B. napus* leaves (petiole-feeding assay) to demonstrate that newly assimilated ^15^N was mostly fixed on “old” ^12^C carbon and that mean ^13^C enrichment of citrate was higher after a light/dark cycle (Gauthier et al., 2010). However, it was not clear to which extent this contribution interfered with the mitochondrial pool of citrate and no quantification of the absolute amount of citrate was done. In addition, a flux balance analysis using a diel model suggested that this stored citrate contribution was explained by the need to support continued export of sugar and amino acids from the leaf during the night and to meet cellular maintenance costs (Cheung et al., 2014). However, previous NMR-based works used both detached leaves to propose the contribution of the stored citrate, which is an experimental setup that necessarily stops sink/source relationships (sugar and amino acids exports) (Gauthier et al., 2010). Here, using *B. napus* leaf discs and a U-^13^C-pyruvate labeling incorporation, we showed that the M2/(M0+M2) ratio of the acetyl-CoA active pool was close to 0.4 after 4 to 6 hours in our experiments. This clearly suggested that most of M0 isotopologues from mitochondrial active pools of TCA cycle metabolites came from M0 acetyl-CoA and M0 oxaloacetate and not a stored pool of ^12^C-citrate (**Figures 6B,8**). Given the important production of PEPc-derived molecules with non-labeled PEP in our experiments, ^12^C-pyruvate necessarily arisen from glycolysis thereby diluting the mitochondrial U-^13^C-pyruvate pool (nearly 20% of M0 still detected for the pyruvate pool at the leaf level). Nevertheless, the concept of stored citrate could also rely on the light-dependent activation of ATP citrate lyase, by converting stored ^12^C-citrate pool into ^12^C-oxaloacetate and ^12^C-acetyl-CoA (Sweetlove et al., 2010;da Fonseca-Pereira et al., 2021). In our experiments the ^13^C-enrichment of acetyl-CoA was indirectly estimated from ^13^C-labeled fragments of either succinate or glutamate, *i.e*. this measurement could not capture a potential contribution of stored ^12^C-citrate to ^12^C-acetyl-CoA. But if such contribution was relatively important in the light and negligible in the dark (concept of the stored citrate contribution), it would necessarily introduce a ^12^C-isotopic dilution for mean ^13^C enrichment of glutamate and succinate in the light compared to the dark (Gauthier et al., 2010). Surprisingly, this isotopic dilution was not observed here while the system was in a metabolic steady-state, thus ruling out a possible bypass with ATP citrate lyase (**Figure 4**). Besides this, we did not observe any significant modification of the citrate pool during the 6 hours of our experiment (**Figure 4B, Table S4**) while a strong decrease in light conditions and a strong increase in dark conditions were expected with the concept of the stored citrate contribution (Cheung et al., 2014). One may question the use of pyruvate instead of CO_2_ (previous works) since pyruvate incorporation may have artificially increased PEPc activity, leading to an important import of malate into mitochondria that stimulate the export of citrate rather than its import through the dicarboxilate carrier 2 (DIC2) (Lee et al., 2021). This could in turn partially explain why the proportion of molecules that experienced a second “turn” of the TCA cycle was relatively weak for citrate, glutamate and succinate compared to malate in our experiments. Nevertheless, it seems more plausible that stored citrate contribution to glutamate biosynthesis was fully decoupled from mitochondrial pools to explain our results (**Figure 8**) (Sweetlove et al., 2010). Overall, our results casted serious doubt about the contribution of stored citrate to the mitochondrial TCA cycle in *B. napus* leaves.

**Figure 8.**
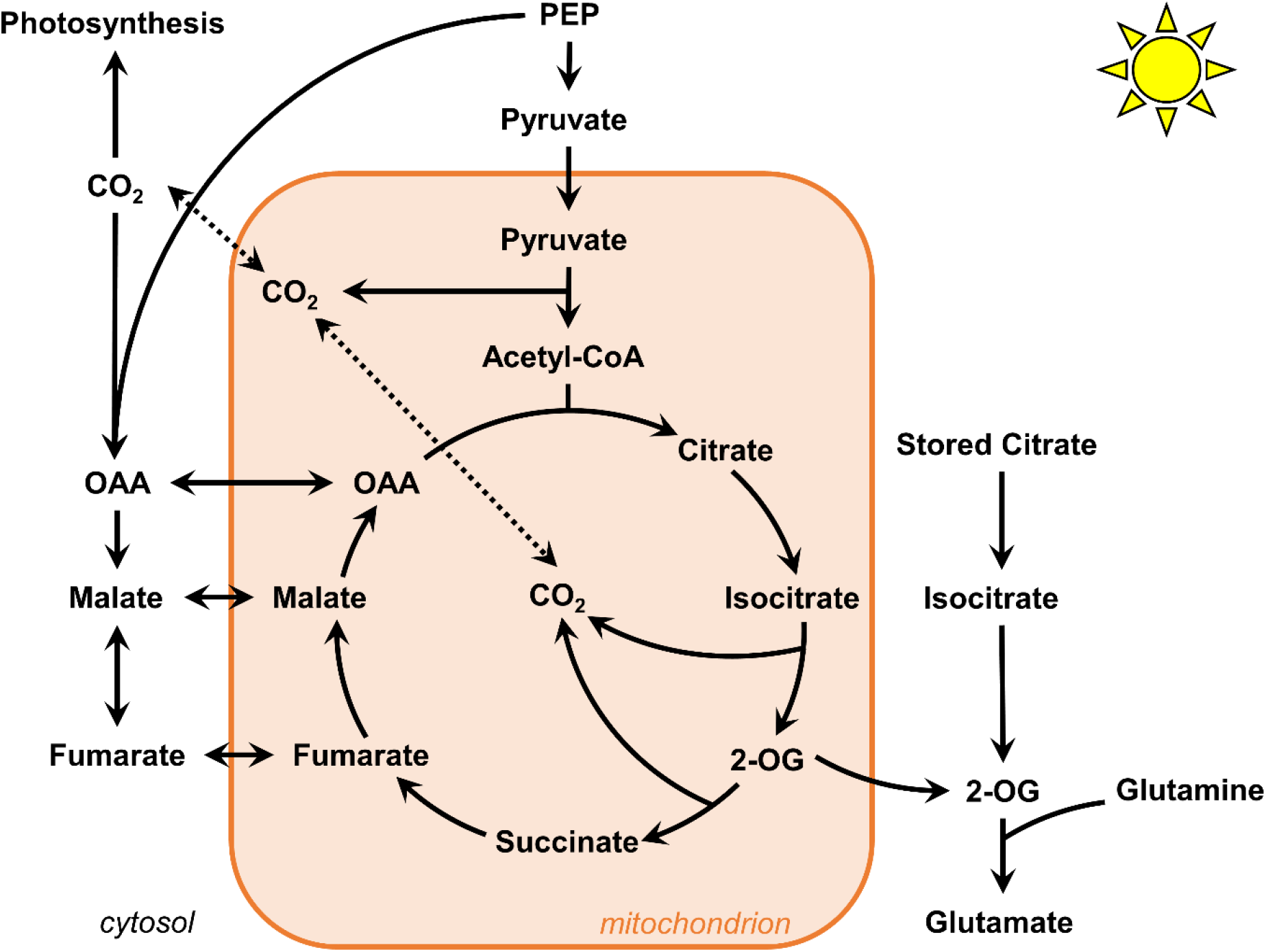
Proposed flux modes for TCA cycle in *B. napus* leaves under light conditions. The contribution of stored citrate is unlikely to interact with mitochondrial TCA cycle. The mitochondrial TCA cycle operate in both cyclic and non-cyclic flux modes. To which extent 2-oxoglutarate molecules escape the cycle and imbalance this cyclic/non-cyclic ratio remain to be determined.

### The TCA cycle partially works in a cyclic mode under light conditions in *B. napus* leaves

The non-cyclic nature of the TCA cycle in illuminated leaves has been experimentally demonstrated from works that used a 1-hour incorporation of ^13^CO_2_ to either detached or attached leaves of different plant species (Tcherkez et al., 2009;Xu et al., 2021;Xu et al., 2022). Only one work used a 2-hour incorporation of ^13^C_1_-pyruvate and ^13^C_3_-pyruvate (Tcherkez et al., 2009), but the major argument for the non-cyclic nature was questionable given that the reassimilation of ^13^CO_2_ arising from ^13^C_1_-pyruvate decarboxylation by PEPc cannot be metabolically transmitted from malate to succinate since the C1 and the C4 of the oxaloacetate backbone are lost during citrate to succinate conversion (**Figure 6A**) (Alves et al., 2015;Lima et al., 2021). Overall, the two INST-MFA approaches showed that PEPc flux and the citrate-to-2-oxoglutarate conversion experienced both relatively similar rates (ratio close to 1), while the flux from 2-oxoglutarate to succinate and from oxaloacetate to malate were both equal to 0 in Arabidopsis and *Camelina sativa* (Xu et al., 2021;Xu et al., 2022). Here, after 2 hours of labeling with U-^13^C-pyruvate, we observed a significant production of TCA cycle-derived isotopologues in citrate, glutamate, succinate and malate fragments followed by an increase of other isotopologues from 2 to 6 hours that corresponded to labeled molecules experiencing a second “turn” of the TCA cycle. Our results provided for the first time in plants a unique ^13^C-labeling signature demonstrating that the TCA cycle can work in a cyclic mode under light conditions in *B. napus* leaves (**Figures 5,6,8**). Therefore, how can we explain discrepancies compared to previous works ? Perhaps, previous works may have missed this cyclic mode due to a shorter time of ^13^C incorporation, but also due to the ^13^C labeling strategy. Indeed, pyruvate can be more rapidly metabolized through the TCA cycle compared to CO_2_. In addition, ^13^CO_2_ could be quickly assimilated by the PEPc thanks to the anhydrase carbonic while the ^13^CO_2_ fixed by the RuBiSCO will require more time to be transmitted to PEP and pyruvate (See the dynamics of CIDs for trioses phosphates, alanine, citrate and aspartate therein (Xu et al., 2021)). In this scenario, ^13^C enrichment of citrate and glutamate will be artificially higher than for succinate (C1 and the C4 of the oxaloacetate backbone are lost during citrate to succinate conversion (**Figure 5A**)). Conversely, it is also important to question our labeling strategy (10 mM U-^13^C-pyruvate incorporations) since it could artificially increase TCA cycle turn-over compared to experiments with ^13^CO_2_ by eliminating the glycolytic regulation step, leading for example to the rapid apparition of M1 and M3 isotopologues for Glutamate_2TMS_C2C5 and Succinate_2TMS_C1C4. A recent incorporation of 15 mM U-^13^C-pyruvate or 15 mM U-^13^C-glucose into Arabidopsis with a petiole-feeding assay during 4 hours in the light identified important ^13^C enrichments in citrate, glutamate and succinate but the CIDs were not shown (Zhang et al., 2021). However, another experiment (Arabidopsis leaf discs, 20 mM U-^13^C-pyruvate, up to 8 hours in the dark) showed similar results but only observed M2 isotopologues for citrate, glutamate and succinate and not the M1 and M3 isotopologues that were observed in our experiments (complete dataset available on request from the authors) (Le et al., 2021). Against these observations for Arabidopsis, it is evident that the important commitment of TCA cycle-derived malate towards another “turn” of the TCA cycle is specific to *B. napus* (**Figure 8**). Therefore, part of the discrepancies between our results and previous works may also be explained by the use of different plant species. Since the non-cyclic nature of the TCA cycle in the light was mostly explained by the need to sustain *de novo* nitrogen assimilation (Gauthier et al., 2010), the analysis of 2-oxoglutarate molecule escaping the mitochondrial TCA cycle should stimulate particular attention in ^13^C labeling experiments in order to compare the rates of cyclic and non-cyclic flux modes (**Figure 8**). For this purpose, the potential role of gamma-aminobutyric acid shunt and glutamate dehydrogenase in rerouting glutamate towards either 2-oxoglutarate or succinate may have to be considered regarding some important studies (Tcherkez et al., 2009;Terce-Laforgue et al., 2013;Nissen et al., 2015).

### Light/dark regulation of PEPc and TCA cycle fluxes in *Brassica napus* leaves

PEPc activity is positively regulated by its light-dependent thioredoxin-mediated phosphorylation in C_3_ and C_4_ plants (Caburatan and Park, 2021). Consequently, PEPc is expected to be more active in the light than in the dark, as recently suggested in guard cell cultures based on the differential ^13^C enrichment found for the C1 of glutamate (Lima et al., 2021). Conversely, TCA cycle is inhibited in light conditions compared to dark conditions, notably due to the inhibition of the PDC and the inhibition of TCA cycle dehydrogenases by the photorespiratory activity of the mitochondrial GDC (Tcherkez et al., 2005;Timm and Hagemann, 2020). Here, we took advantage of pathway specific isotopologue/isotopomer to calculate fractional contribution (FC) of TCA cycle and PEPc for pyruvate-to-malate and pyruvate-to-citrate. We showed that the contribution of PEPc-dependent reassimilation of TCA cycle and PDC-derived CO_2_ was relatively low compared to the TCA cycle for malate (4:1 ratio) but was positively regulated by the light condition (1.5-fold increase), reflecting the strong shift for M1/M2 isotopologues of Malate_3TMS_C1C4 in light conditions compared to dark conditions (**Figures 6,8**). Considering the relative nature of pathway FC, this shift could be attributed to both increased PEPc activity and TCA cycle inhibition in the light. However, in our experiments, we found very similar dynamics of ^13^C-enrichment for TCA cycle metabolites in both dark and light conditions combined to similar metabolic pool sizes in a metabolic steady-state (**Figure 4**). Thus, variations of TCA cycle metabolic fluxes between light and dark conditions are expected to be very low. This may be attributed to the artificial incorporation of pyruvate that eliminated the PDC-dependent regulation of carbon entry for plant TCA cycle. Nevertheless, the importance of this regulation for carbon entry towards the TCA cycle can also be questioned. A previous work based on ^13^CO_2_ decarboxylations with either ^13^C-pyruvate or ^13^C-glucose showed that PDC was inhibited by only 27% in the light compared to the dark, while glycolysis could be inhibited by up to 95% (Tcherkez et al., 2005). Recently, a U-^13^C-pyruvate incorporation into Arabidopsis leaves during 4 hours in the light showed that the *pdk* mutant (constant activation of the PDC) had higher ^13^C-enrichments for citrate, glutamate and succinate compared to the control but these differences were not observed when using U-^13^C-glucose (Zhang et al., 2021). Based on these observations, we think that pyruvate incorporation essentially eliminated the light/dark regulation of glycolysis in our experiments. Thus, our ^13^C-labeling strategy can only be used to probe light/dark regulation of PEPc activity, through the reassimilation of TCA cycle and PDC-derived CO_2_. To which extent this PEPc-dependent reassimilation is close to the total PEPc flux remains an important question to address for a complete comparison and TCA cycle and PEPc fluxes. For instance, the two ^13^C-INST-MFA performed in plant leaves with ^13^CO_2_ obtained a 1:1 ratio with only non-cyclic flux mode for the TCA cycle. A^13^C-MFA performed at isotopic and metabolic steady-state in heterotrophic suspension cells culture incubated with U-^13^C-glucose showed a similar ratio of 2:1 for forward TCA cycle/PEPc metabolic fluxes (ana1 versus tca2, tca3, tca4, tca5), with a negligible malic enzyme activity (10-fold less than PEPc) (Masakapalli et al., 2010). Whether the total PEPc flux is higher than the cyclic mode of the TCA cycle for both malate and citrate in *B. napus* leaves will have to be addressed using complementary ^13^C-labeling strategy, such U-^13^C-glucose or U-^13^C-sucrose, or with a full ^13^CO_2_-based INST-MFA with longer periods of labeling compared to previous works. Besides this, we observed an unexpected higher contribution of PEPc derived-carbons to citrate in the dark condition compared to the light condition. Since we have shown that PEPc contribution to malate was reduced in the dark compared to the light, this result could reflect a higher transport of oxaloacetate/malate from the cytosol to the mitochondria in dark conditions. This idea is supported by the fact that PEPc is majorly localized to the cytosol (Caburatan and Park, 2021) and the recent identification of a mitochondrial malate importer in Arabidopsis with an important impact on mitochondrial respiration in the dark (Lee et al., 2021). The import/export of malate and citrate in mitochondria should also be considered in future ^13^C-MFA of the TCA cycle.

## Conclusions

Overall, we proposed a robust workflow to evaluate the accuracy of CID measurements by GC-MS for organic and amino acids and successfully applied our validated method to the investigation of TCA cycle and PEPc metabolic fluxes in *Brassica napus* leaves using a ^13^C-MFA concept. Our results did not support the hypothesis of the stored citrate contribution to mitochondrial TCA cycle in a light/dark-dependent manner and demonstrated that the forward cyclic flux mode of the TCA cycle can operate under light conditions, leading to nonnegligible commitment of *de novo* biosynthesized TCA cycle intermediates to another “turn” of the cycle (**Figure 8**). We also showed that pathway fractional contribution could be used to probe the light/dark regulation of PEPc-dependent reassimilation of mitochondrial decarboxylations. Notably, the reassimilation of PDC and TCA-cycle derived CO_2_ by photosynthesis represented nearly 5% of the net CO_2_ assimilation rate in our conditions. The next stage of this work could be focused on the accurate modeling of this metabolic network to infer absolute metabolic fluxes. This perspective sounds stimulating given the development of multiple modeling approaches over the last decade but still remains highly challenging (Kruger and Ratcliffe, 2021).

## 6 Author Contributions

Conceptualization, Y.D., A.B.; methodology, Y.D..; software, Y.D., O.F.; validation, Y.D., S.B. and C.B.; formal analysis, Y.D.; investigation, Y.D., S.B. and C.B.; data curation, Y.D.; writing-original draft preparation, Y.D.; writing-review and editing, Y.D., A.B.; visualization, Y.D.; supervision, Y.D.; project administration, Y.D.; funding acquisition, Y.D., A.B. All authors have read and agreed to the published version of the manuscript.

## 8 Acknowledgments

P2M2 (Metabolic Profiling & Metabolomics Platform, Le Rheu, France, www6.inrae.fr/p2m2/) and its staff members are gratefully acknowledged for access to mass spectrometry facilities and for supporting the methodological developments required for this study. We thank MetaToul (Metabolomics & Fluxomics Facitilies, Toulouse, France, www.metatoul.fr) for technical support and access to NMR facilities. We are grateful to Nathalie Marnet and Mathieu Aubert for technical assistance in the quantification of amino acid content and Pierre Millard and Floriant Bellvert for fruitful scientific discussions.

## 9 Supplementary Material

Supplementary Material is available upon request (issues with uploading on the biorxiv server).

## References

Abadie, C., Lothier, J., Boex-Fontvieille, E., Carroll, A., and Tcherkez, G. (2017). Direct assessment of the metabolic origin of carbon atoms in glutamate from illuminated leaves using (13) C-NMR. New Phytol 216, 1079–1089.

Abadie, C., and Tcherkez, G. (2019). In vivo phosphoenolpyruvate carboxylase activity is controlled by CO2 and O2 mole fractions and represents a major flux at high photorespiration rates. New Phytol 221, 1843–1852.

Abadie, C., and Tcherkez, G. (2021). (13)C Isotope Labelling to Follow the Flux of Photorespiratory Intermediates. Plants (Basel) 10.

Allen, D.K., and Young, J.D. (2019). Tracing metabolic flux through time and space with isotope labeling experiments. Curr Opin Biotechnol 64, 92–100.

Alves, T.C., Pongratz, R.L., Zhao, X., Yarborough, O., Sereda, S., Shirihai, O., Cline, G.W., Mason, G., and Kibbey, R.G. (2015). Integrated, Step-Wise, Mass-Isotopomeric Flux Analysis of the TCA Cycle. Cell Metab 22, 936–947.

Antoniewicz, M.R. (2021). A guide to metabolic flux analysis in metabolic engineering: Methods, tools and applications. Metab Eng 63, 2–12.

Antoniewicz, M.R., Kelleher, J.K., and Stephanopoulos, G. (2006). Determination of confidence intervals of metabolic fluxes estimated from stable isotope measurements. Metab Eng 8, 324–337.

Antoniewicz, M.R., Kelleher, J.K., and Stephanopoulos, G. (2007). Accurate Assessment of Amino Acid Mass Isotopomer Distributions for Metabolic Flux Analysis. Anal Chem 79, 6.

Araujo, W.L., Nunes-Nesi, A., Nikoloski, Z., Sweetlove, L.J., and Fernie, A.R. (2012). Metabolic control and regulation of the tricarboxylic acid cycle in photosynthetic and heterotrophic plant tissues. Plant Cell Environ 35, 1–21.

Araujo, W.L., Tohge, T., Nunes Nesi, A., Obata, T., and Fernie, A. (2014). “Analysis of Kinetic Labeling of Amino Acids and Organic Acids by GC-MS,” in Plant Metabolic Flux Analysis: Methods and Protocols, ed. M.I.M. Biology.).

Bianchetti, G., Baron, C., Carrillo, A., Berardocco, S., Marnet, N., Wagner, M.H., Demilly, D., Ducournau, S., Manzanares-Dauleux, M.J., Caherec, F.L., Buitink, J., and Nesi, N. (2021). Dataset for the metabolic and physiological characterization of seeds from oilseed rape (Brassica napus L.) plants grown under single or combined effects of drought and clubroot pathogen Plasmodiophora brassicae. Data Brief 37, 107247.

Buescher, J.M., Antoniewicz, M.R., Boros, L.G., Burgess, S.C., Brunengraber, H., Clish, C.B., Deberardinis, R.J., Feron, O., Frezza, C., Ghesquiere, B., Gottlieb, E., Hiller, K., Jones, R.G., Kamphorst, J.J., Kibbey, R.G., Kimmelman, A.C., Locasale, J.W., Lunt, S.Y., Maddocks, O.D., Malloy, C., Metallo, C.M., Meuillet, E.J., Munger, J., Noh, K., Rabinowitz, J.D., Ralser, M., Sauer, U., Stephanopoulos, G., St-Pierre, J., Tennant, D.A., Wittmann, C., Vander Heiden, M.G., Vazquez, A., Vousden, K., Young, J.D., Zamboni, N., and Fendt, S.M. (2015). A roadmap for interpreting (13)C metabolite labeling patterns from cells. Curr Opin Biotechnol 34, 189–201.

Caburatan, L., and Park, J. (2021). Differential Expression, Tissue-Specific Distribution, and Posttranslational Controls of Phosphoenolpyruvate Carboxylase. Plants (Basel) 10.

Cheah, Y.E., Xu, Y., Sacco, S.A., Babele, P.K., Zheng, A.O., Johnson, C.H., and Young, J.D. (2020). Systematic identification and elimination of flux bottlenecks in the aldehyde production pathway of Synechococcus elongatus PCC 7942. Metab Eng 60, 56–65.

Cheah, Y.E., and Young, J.D. (2018). Isotopically nonstationary metabolic flux analysis (INST-MFA): putting theory into practice. Curr Opin Biotechnol 54, 80–87.

Cheung, C.Y., Poolman, M.G., Fell, D.A., Ratcliffe, R.G., and Sweetlove, L.J. (2014). A Diel Flux Balance Model Captures Interactions between Light and Dark Metabolism during Day-Night Cycles in C3 and Crassulacean Acid Metabolism Leaves. Plant Physiol 165, 917–929.

Chu, K.L., Koley, S., Jenkins, L.M., Bailey, S.R., Kambhampati, S., Foley, K., Arp, J.J., Morley, S.A., Czymmek, K.J., Bates, P.D., and Allen, D.K. (2022). Metabolic flux analysis of the non-transitory starch tradeoff for lipid production in mature tobacco leaves. Metab Eng 69, 231–248.

Clark, T.J., Guo, L., Morgan, J., and Schwender, J. (2020). Modeling Plant Metabolism: From Network Reconstruction to Mechanistic Models. Annu Rev Plant Biol.

Da Fonseca-Pereira, P., Souza, P.V.L., Fernie, A.R., Timm, S., Daloso, D.M., and Araujo, W.L. (2021). Thioredoxin-mediated regulation of (photo)respiration and central metabolism. J Exp Bot 72, 5987–6002.

Daloso, D.M., Medeiros, D.B., Dos Anjos, L., Yoshida, T., Araujo, W.L., and Fernie, A.R. (2017). Metabolism within the specialized guard cells of plants. New Phytol 216, 1018–1033.

Dellero, Y. (2020). Manipulating amino acid metabolism to improve crop nitrogen use efficiency for a sustainable agriculture. Frontiers in Plant Science 11, 1857

Dellero, Y., Clouet, V., Marnet, N., Pellizzaro, A., Dechaumet, S., Niogret, M.F., and Bouchereau, A. (2020a). Leaf status and environmental signals jointly regulate proline metabolism in winter oilseed rape. J Exp Bot 71, 2098–2111.

Dellero, Y., and Filangi, O. (2021). “Corrective method dedicated to Isocor for calculating carbon isotopologue distribution from GCMS runs”. V1 ed.: Portail Data INRAE).

Dellero, Y., Heuillet, M., Marnet, N., Bellvert, F., Millard, P., and Bouchereau, A. (2020b). Sink/Source Balance of Leaves Influences Amino Acid Pools and Their Associated Metabolic Fluxes in Winter Oilseed Rape (Brassica napus L.). Metabolites 10, 16.

Dellero, Y., Jossier, M., Glab, N., Oury, C., Tcherkez, G., and Hodges, M. (2016). Decreased glycolate oxidase activity leads to altered carbon allocation and leaf senescence after a transfer from high CO2 to ambient air in Arabidopsis thaliana. J Exp Bot 67, 3149–3163.

Des Rosiers, C., Lloyd, S., Comte, B., and Chatham, J.C. (2004). A critical perspective of the use of (13)C-isotopomer analysis by GCMS and NMR as applied to cardiac metabolism. Metab Eng 6, 44–58.

Dong, H., Bai, L., Zhang, Y., Zhang, G., Mao, Y., Min, L., Xiang, F., Qian, D., Zhu, X., and Song, C.P. (2018). Modulation of Guard Cell Turgor and Drought Tolerance by a Peroxisomal Acetate-Malate Shunt. Mol Plant 11, 1278–1291.

Evers, B., Gerding, A., Boer, T., Heiner-Fokkema, M.R., Jalving, M., Wahl, S.A., Reijngoud, D.J., and Bakker, B.M. (2021). Simultaneous Quantification of the Concentration and Carbon Isotopologue Distribution of Polar Metabolites in a Single Analysis by Gas Chromatography and Mass Spectrometry. Anal Chem 93, 8248–8256.

Galili, G., Amir, R., and Fernie, A.R. (2016). The Regulation of Essential Amino Acid Synthesis and Accumulation in Plants. Annu Rev Plant Biol 67, 153–178.

Gauthier, P.P., Bligny, R., Gout, E., Mahe, A., Nogues, S., Hodges, M., and Tcherkez, G.G. (2010). In folio isotopic tracing demonstrates that nitrogen assimilation into glutamate is mostly independent from current CO2 assimilation in illuminated leaves of Brassica napus. New Phytol 185, 988–999.

Harvey, D.J., and Vouros, P. (2020). Mass Spectrometric Fragmentation of Trimethylsilyl and Related Alkylsilyl Derivatives. Mass Spectrom Rev 39, 105–211.

Heuillet, M., Bellvert, F., Cahoreau, E., Letisse, F., Millard, P., and Portais, J.C. (2018). Methodology for the Validation of Isotopic Analyses by Mass Spectrometry in Stable-Isotope Labeling Experiments. Anal Chem 90, 1852–1860.

Heuillet, M., Millard, P., Cisse, M.Y., Linares, L.K., Letisse, F., Manie, S., Le Cam, L., Portais, J.C., and Bellvert, F. (2020). Simultaneous Measurement of Metabolite Concentration and Isotope Incorporation by Mass Spectrometry. Anal Chem.

Kambhampati, S., Li, J., Evans, B.S., and Allen, D.K. (2019). Accurate and efficient amino acid analysis for protein quantification using hydrophilic interaction chromatography coupled tandem mass spectrometry. Plant Methods 15, 46.

Kruger, N.J., Huddleston, J.E., Le Lay, P., Brown, N.D., and Ratcliffe, R.G. (2007). Network flux analysis: impact of 13C-substrates on metabolism in Arabidopsis thaliana cell suspension cultures. Phytochemistry 68, 2176–2188.

Kruger, N.J., and Ratcliffe, R.G. (2021). Whither metabolic flux analysis in plants? Journal of Experimental Botany.

Lai, Z., and Fiehn, O. (2018). Mass spectral fragmentation of trimethylsilylated small molecules. Mass Spectrom Rev 37, 245–257.

Le, X.H., Lee, C.P., and Millar, A.H. (2021). The mitochondrial pyruvate carrier (MPC) complex mediates one of three pyruvate-supplying pathways that sustain Arabidopsis respiratory metabolism. Plant Cell 33, 2776–2793.

Le, X.H., Lee, C.P., Monachello, D., and Millar, A.H. (2022). Metabolic evidence for distinct pyruvate pools inside plant mitochondria. Nat Plants.

Lee, C.P., Elsasser, M., Fuchs, P., Fenske, R., Schwarzlander, M., and Millar, A.H. (2021). The versatility of plant organic acid metabolism in leaves is underpinned by mitochondrial malate-citrate exchange. Plant Cell 33, 3700–3720.

Lima, V.F., Erban, A., Daubermann, A.G., Freire, F.B.S., Porto, N.P., Candido-Sobrinho, S.A., Medeiros, D.B., Schwarzlander, M., Fernie, A.R., Dos Anjos, L., Kopka, J., and Daloso, D.M. (2021). Establishment of a GC-MS-based (13) C-positional isotopomer approach suitable for investigating metabolic fluxes in plant primary metabolism. Plant J.

Lima, V.F., Perez Souza, L., Williams, T.C., Fernie, A., and Daloso, D.M. (2018). “Gas Chromatography–Mass Spectrometry-Based 13C-Labeling Studies in Plant Metabolomics,” in Plant Metabolomics: Methods and Protocol, ed. M.I.M. Biology.).

Ma, F., Jazmin, L.J., Young, J.D., and Allen, D.K. (2014). Isotopically nonstationary 13C flux analysis of changes in Arabidopsis thaliana leaf metabolism due to high light acclimation. Proc Natl Acad Sci U S A 111, 16967–16972.

Masakapalli, S.K., Bryant, F.M., Kruger, N.J., and Ratcliffe, R.G. (2014). The metabolic flux phenotype of heterotrophic Arabidopsis cells reveals a flexible balance between the cytosolic and plastidic contributions to carbohydrate oxidation in response to phosphate limitation. Plant J 78, 964–977.

Masakapalli, S.K., Le Lay, P., Huddleston, J.E., Pollock, N.L., Kruger, N.J., and Ratcliffe, R.G. (2010). Subcellular flux analysis of central metabolism in a heterotrophic Arabidopsis cell suspension using steady-state stable isotope labeling. Plant Physiol 152, 602–619.

Medeiros, D.B., Perez Souza, L., Antunes, W.C., Araujo, W.L., Daloso, D.M., and Fernie, A.R. (2018). Sucrose breakdown within guard cells provides substrates for glycolysis and glutamine biosynthesis during light-induced stomatal opening. Plant J 94, 583–594.

Millard, P., Cahoreau, E., Heuillet, M., Portais, J.C., and Lippens, G. (2017). (15)N-NMR-Based Approach for Amino Acids-Based (13)C-Metabolic Flux Analysis of Metabolism. Anal Chem 89, 2101–2106.

Millard, P., Delepine, B., Guionnet, M., Heuillet, M., Bellvert, F., and Letisse, F. (2019). IsoCor: isotope correction for high-resolution MS labeling experiments. Bioinformatics 35, 4484–4487.

Millard, P., Massou, S., Portais, J.C., and Letisse, F. (2014). Isotopic studies of metabolic systems by mass spectrometry: using Pascal’s triangle to produce biological standards with fully controlled labeling patterns. Anal Chem 86, 10288–10295.

Millard, P., Schmitt, U., Kiefer, P., Vorholt, J.A., Heux, S., and Portais, J.-C. (2020). ScalaFlux: a scalable approach to quantify fluxes in metabolic subnetworks. Plos Computational Biology 16, e1007799.

Nissen, J.D., Pajecka, K., Stridh, M.H., Skytt, D.M., and Waagepetersen, H.S. (2015). Dysfunctional TCA-Cycle Metabolism in Glutamate Dehydrogenase Deficient Astrocytes. Glia 63, 2313–2326.

Okahashi, N., Kawana, S., Iida, J., Shimizu, H., and Matsuda, F. (2019). Fragmentation of Dicarboxylic and Tricarboxylic Acids in the Krebs Cycle Using GC-EI-MS and GC-EI-MS/MS. Mass Spectrom (Tokyo) 8, A0073.

Ratcliffe, R.G., and Shachar-Hill, Y. (2006). Measuring multiple fluxes through plant metabolic networks. Plant J 45, 490–511.

Rstudio Team (2016). RStudio: Integrated Development for R.

Sweetlove, L.J., Beard, K.F., Nunes-Nesi, A., Fernie, A.R., and Ratcliffe, R.G. (2010). Not just a circle: flux modes in the plant TCA cycle. Trends Plant Sci 15, 462–470.

Szecowka, M., Heise, R., Tohge, T., Nunes-Nesi, A., Vosloh, D., Huege, J., Feil, R., Lunn, J., Nikoloski, Z., Stitt, M., Fernie, A.R., and Arrivault, S. (2013). Metabolic fluxes in an illuminated Arabidopsis rosette. Plant Cell 25, 694–714.

Tcherkez, G., Cornic, G., Bligny, R., Gout, E., and Ghashghaie, J. (2005). In vivo respiratory metabolism of illuminated leaves. Plant Physiol 138, 1596–1606.

Tcherkez, G., Gauthier, P., Buckley, T.N., Busch, F.A., Barbour, M.M., Bruhn, D., Heskel, M.A., Gong, X.Y., Crous, K.Y., Griffin, K., Way, D., Turnbull, M., Adams, M.A., Atkin, O.K., Farquhar, G.D., and Cornic, G. (2017). Leaf day respiration: low CO2 flux but high significance for metabolism and carbon balance. New Phytol 216, 986–1001.

Tcherkez, G., Mahe, A., Gauthier, P., Mauve, C., Gout, E., Bligny, R., Cornic, G., and Hodges, M. (2009). In folio respiratory fluxomics revealed by 13C isotopic labeling and H/D isotope effects highlight the noncyclic nature of the tricarboxylic acid “cycle” in illuminated leaves. Plant Physiol 151, 620–630.

Terce-Laforgue, T., Bedu, M., Dargel-Grafin, C., Dubois, F., Gibon, Y., Restivo, F.M., and Hirel, B. (2013). Resolving the role of plant glutamate dehydrogenase: II. Physiological characterization of plants overexpressing the two enzyme subunits individually or simultaneously. Plant Cell Physiol 54, 1635–1647.

Timm, S. (2020). The impact of photorespiration on plant primary metabolism through metabolic and redox regulation. Biochemical Society Transactions 48, 2495–2504.

Timm, S., and Hagemann, M. (2020). Photorespiration - how is it regulated and regulates overall plant metabolism? J Exp Bot.

Treves, H., Kuken, A., Arrivault, S., Ishihara, H., Hoppe, I., Erban, A., Hohne, M., Moraes, T.A., Kopka, J., Szymanski, J., Nikoloski, Z., and Stitt, M. (2022). Carbon flux through photosynthesis and central carbon metabolism show distinct patterns between algae, C3 and C4 plants. Nat Plants 8, 78–91.

Xu, G., Fan, X., and Miller, A.J. (2012). Plant nitrogen assimilation and use efficiency. Annu Rev Plant Biol 63, 153–182.

Xu, Y., Fu, X., Sharkey, T.D., Shachar-Hill, Y., and Walker, B.J. (2021). The metabolic origins of non-photorespiratory CO2 release during photosynthesis: A metabolic flux analysis. Plant Physiol.

Xu, Y., Wieloch, T., Kaste, J.a.M., Shachar-Hill, Y., and Sharkey, T.D. (2022). Reimport of carbon from cytosolic and vacuolar sugar pools into the Calvin–Benson cycle explains photosynthesis labeling anomalies. Proceedings of the National Academy of Sciences 119.

Young, J.D., Allen, D.K., and Morgan, J.A. (2014). “Isotopomer measurement techniques in metabolic flux analysis II: mass spectrometry,” in Methods Mol Biol.), 85–108.

Zhang, Y., and Fernie, A.R. (2018). On the role of the tricarboxylic acid cycle in plant productivity. J Integr Plant Biol 60, 1199–1216.

Zhang, Y., Giese, J., Kerbler, S.M., Siemiatkowska, B., Perez De Souza, L., Alpers, J., Medeiros, D.B., Hincha, D.K., Daloso, D.M., Stitt, M., Finkemeier, I., and Fernie, A.R. (2021). Two mitochondrial phosphatases, PP2c63 and Sal2, are required for posttranslational regulation of the TCA cycle in Arabidopsis. Mol Plant 14, 1104–1118.

